# Multivariate AND-gate substrate probes as enhanced contrast agents for fluorescence-guided surgery

**DOI:** 10.1101/695403

**Authors:** John C. Widen, Martina Tholen, Joshua J. Yim, Alexander Antaris, Kerriann M. Casey, Stephan Rogalla, Alwin Klaassen, Jonathan Sorger, Matthew Bogyo

## Abstract

The greatest challenges for surgical management of cancer are precisely locating lesions and clearly defining the margins between tumors and normal tissues. This is confounded by the characteristics of the tissue where the tumor is located as well as its propensity to form irregular boundaries with healthy tissues. To address these issues, molecularly targeted optical contrast agents have been developed to define margins in real-time during surgery^1,2^. However, selectivity of a contrast agent is often limited by expression of a target enzyme or receptor in both tumor and healthy tissues. Here we introduce a concept of multivariate ‘AND-gate’ optical imaging probes that require sequential processing by multiple tumor-specific enzymes to produce a fluorescent signal. This results in dramatically improved specificity as well as overall enhanced sensitivity. This general approach has the potential to be broadly applied to selectively target complex patterns of enzyme activities in diverse disease tissues for detection, treatment and therapy response monitoring.

## Introduction

For many cancers, surgery is the primary treatment option followed by chemo and/or radiation therapy. Early detection and surgical removal of solid tumors currently remain the most effective way to produce a curative result. Success outcomes are highly dependent on how effectively the tumor tissue can be identified^2,3^. Accurate detection of the ‘margin’ between tumor and normal healthy tissues is essential to prevent either incomplete removal of tumor cells leading to increased recurrence rates (up to 30-65% of cases)^4-6^ or removal of excess healthy tissue. Furthermore, solid tumors derived from different tissues and cell types can have dramatically different margin borders, making the prospects for complete removal highly variable^7^. In fact, rates of repeat surgeries for some cancer types (i.e. breast) can be as high as 50%^3,8^. Therefore, new methods that allow both selective and sensitive real-time detection of tumor margins have the potential to significantly impact surgical treatment outcomes of many of the most common types of cancer.

For most surgical treatments for cancer, imaging methods such as MRI, PET, SPECT and CT are used to diagnose and then identify the location of tumors^9^. However, these methods are not easily used intra-operatively due to the need for large instrumentation and/or ionizing radiation. They also typically do not provide sufficient sensitivity and resolution to allow identification of microscopic processes at the tumor margin. Alternatively, various analytical methods can be used to detect specific molecular signatures in real-time during a surgical procedure. This includes the use of mass spectrometry (MS), radio frequency (RF), ultrasound and fluorescence lifetime detectors that allow measurement of specific analytical signatures in the tumor microenvironment that set it apart from the surrounding normal tissues^10-12^. However, while these types of real-time analytical devices can be highly sensitive and accurate, they all are limited by issues related to sampling error and slow processing times since they require scanning of regions suspected to contain residual cancer.

As an alternative to the use of analytical detectors to scan tissues, systemically delivered optical contrast agents have the potential to broadly highlight tumor sites without any prior knowledge of tumor location. Currently, there are only a few optical agents that are FDA approved for use in humans and that are suitable for use with clinical camera systems. While there has been significant clinical benefit from dyes such as ICG^13^ and 5-ALA^14^, the main limitation is their overall non-specific mechanism of action resulting in insufficient sensitivity and selectivity for use in diverse cancer types. A number of pre-clinical and early-stage clinical trials have begun for fluorescently tagged affinity agents such as antibodies, peptides and small molecule ligands^1,2,15^. There have also been significant advances in the development of “smart” probes that only produce a fluorescent signal when processed by a tumor-specific enzyme (**Fig. 1a**). This leads to a further increase in contrast because free probe outside the tumor remains unprocessed and is not fluorescent. Proteases have become the enzymes of choice for generation of smart probes because short peptide substrates can be decorated with suitable fluorophore/quencher pairs such that probes are optically silent until cleaved by a protease^16-19^. Protease activity can also be used to process probes that then change their localization to be retained in cells within the tumor microenvironment^18,20^. Multiple proteases have proven to be useful biomarkers of cancerous tissues^21-24^.

**Figure 1.**
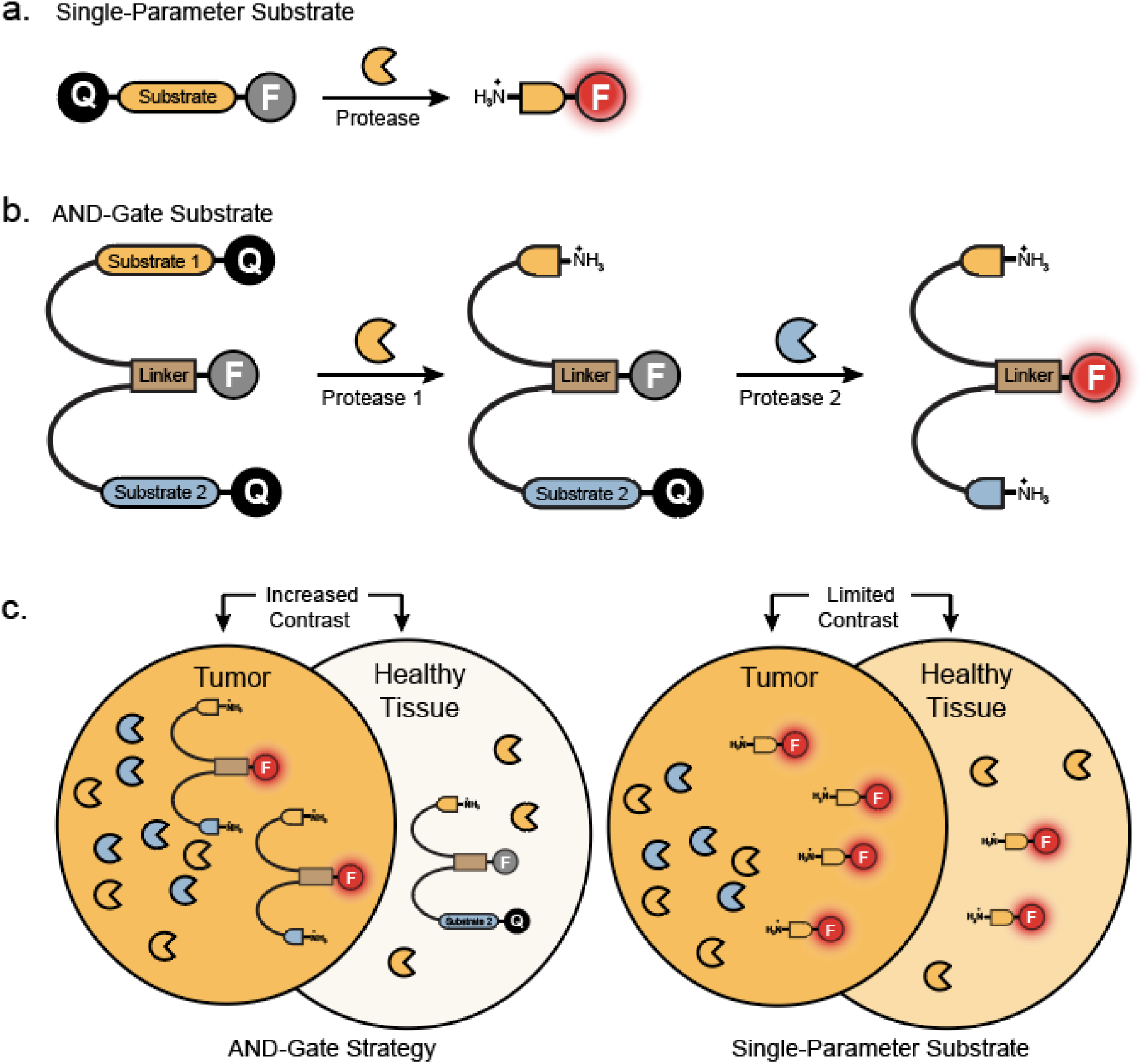
The AND-Gate strategy. (**a**) Schematic of a single-parameter quenched-fluorescent protease substrate that is cleaved by a single protease to produce a fluorescent fragment. These probes take advantage of higher proteolytic activity in tumors to generate contrast. (**b**) Schematic of the multivariate AND-Gate probe, which requires two proteolytic processing events to activate fluorescence. (**c**) Comparison of the single-parameter and AND-Gate probes in tumor and normal tissues. Higher contrast is generated by the AND-gate probe due to lack of activation in normal tissues which lack both protease activities.

Previously, we designed a cysteine cathepsin-cleavable NIR substrate probe 6QC-NIR^20^. Cleavage of this substrate probe produces high signals in cancers of the lung, breast, and colon, which can be detected with the FDA-approved da Vinci^®^ Si Surgical System equipped with the Firefly^®^ detection system^20^. We have found that further optimization of the reporter dye on this probe can greatly enhance its use with clinical camera systems^25^. However, the single biggest limitation to the use of protease substrates is their lack of selectivity that results from activity of a protease or protease family in healthy tissues as well as inside the tumor microenvironment. Here we describe a way to dramatically increase the overall tumor-selectivity by designing multivariate ‘AND-gate’ probes in which multiple reporters must be sequentially processed within the tumor microenvironment to produce a specific optical signal (**Fig. 1b, c**). Thus, selectivity can be greatly enhanced because identification of the tumor tissue is made using multiple markers that help to distinguish it from the surrounding normal tissues. In the current study, we demonstrate the utility of the approach using a probe that contains substrates for both the lysosomal cysteine cathepsins found in tumor associated macrophages and caspases that are activated only in apoptotic cells. The resulting AND-gate probes have greatly improved tumor selectivity due to the relatively high levels of apoptosis in tumors compared to surrounding normal healthy tissues. In addition, we find that AND-gate probes have increased signal accumulation compared to the corresponding cathepsin-only probe. These improved properties of the AND-gate probes suggest that this will be a general strategy that can be used with diverse enzyme substrates to increase the selectivity of optical contrast agents.

## Results

### Design, testing and optimization of AND-Gate Probes

The design of a general ‘AND-gate’ strategy involves placement of a fluorescent reporter in a central location such that multiple quenchers can be attached through linkages that are severed by the action of enzymes. This type of ‘hub and spoke’ model requires removal of all quenchers along the spokes to liberate the central fluorescent hub. In theory, the quencher groups can be attached to the central hub using any type of enzyme-sensitive linkage resulting in AND-gates that are responsive to a diverse array of enzyme signatures. We choose to test this design strategy using a simple glutamic acid (Glu) central linker containing a fluorescent dye linked to quenchers through distinct peptide sequences that could be orthogonally cleaved by two proteases with non-overlapping substrate specificities. The central Glu linker is ideal because a sulfo-Cy5 fluorophore can be attached to its free α-amine and each peptide substrate containing a sulfo-QSY21 quencher can be connected through amide bonds to the remaining carboxylic acids (**Fig. 2a**). This positions the quenchers such that fluorescence activation occurs only after both peptides have been cleaved. Furthermore, by using diamino-alkyl linkers between the peptides and the Glu core, the final fluorescent fragment will contain two free amine groups that induce lysosomal accumulation of the probe, as we have demonstrated for our cathepsin substrate probes^20^. We chose peptide substrates that have already been shown to be specifically processed by caspase 3 (Casp3; DEVDGP)^26^ and the cysteine cathepsins (Cats; Z-FK)^27^. These sequences are optimal for a first generation AND-gate probe because Casp3 and Cats have highly orthogonal substrate specificities and therefore each protease can only process its corresponding substrate. Furthermore, Cats are upregulated in both tumor associated macrophages (TAMs)^28-30^ and in most normal tissues, while Casp3 activity is elevated in tumors but is generally not active in healthy cells, except during development^31^. Therefore, Cats and Casp3 are both active within tumors but are not expected to be active together in healthy tissues, making them ideal targets for our AND-gate probe strategy.

**Figure 2.**
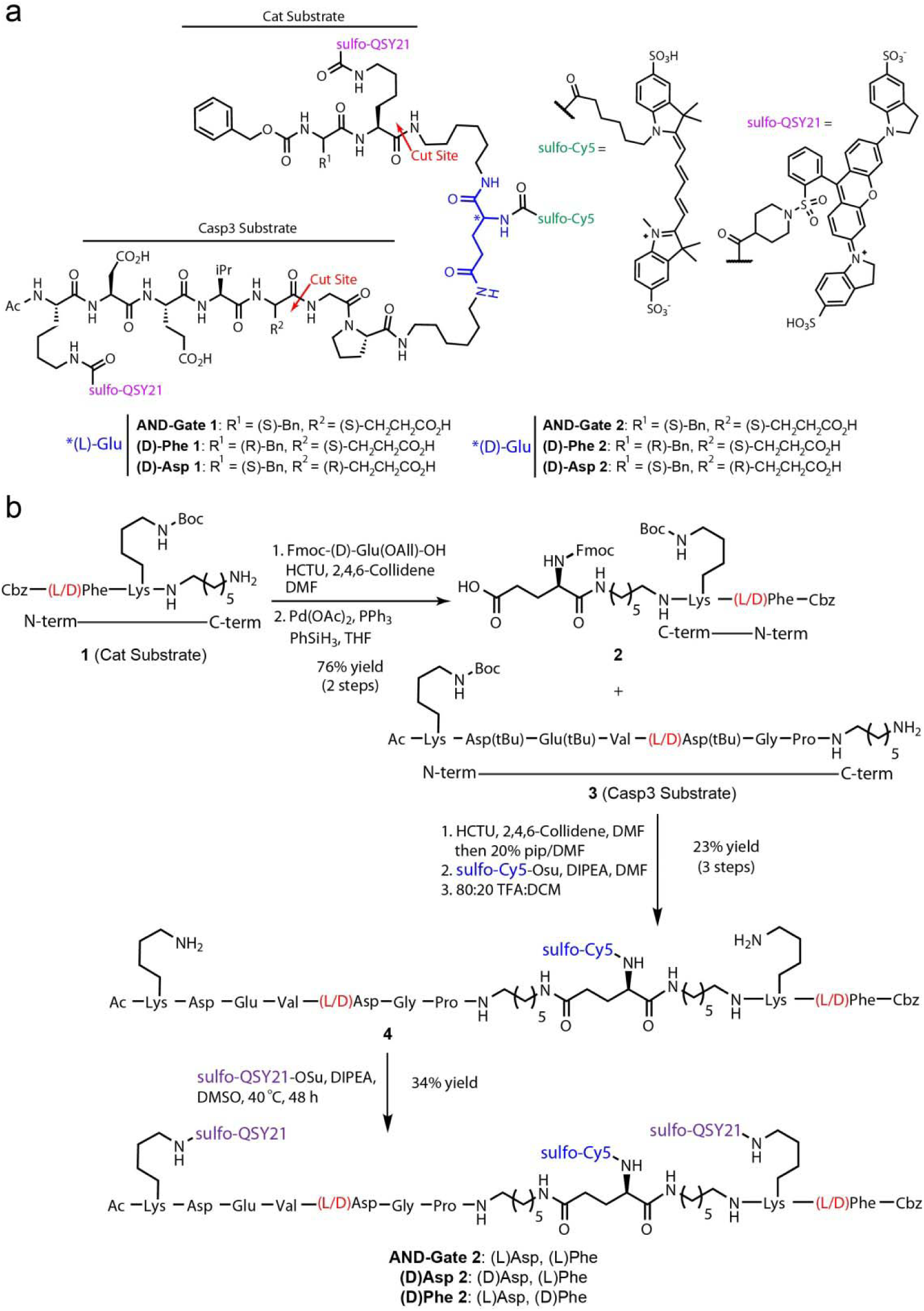
Design and synthesis of AND-gate probes. (a) Structures of the dual orthogonal protease substrate AND-Gate probes and negative controls. **AND-Gate 1** contains the natural amino acid (L)-Glu while the **AND-Gate 2** contains a non-natural (D)-Glu linker (blue). The negative control probes contain either a (D)-Asp in the P1 position of the Casp3 sequence (**(D)-Asp 1**) or (D)-Phe in the P2 position of the Cat sequence (**(D)-Phe 1**) to block proteolytic cleavage by the respective protease. (b) Synthesis of **AND-Gate 2, (D)Asp 2**, and **(D)Phe 2** containing a (D)-Glu central linker. See Methods section for additional synthetic details.

We synthesized a first-generation probe, **AND-Gate 1**, containing the natural (L)-Glu central linker (**Fig. 1d, Supplementary Scheme 1**). We also synthesized negative controls in which a (D)-Asp was used in place of the natural (L)-Asp in the P1 position of the Casp3 sequence (**(D)-Asp 1**) and a (D)-Phe was used in the P2 position of the Cat sequence (**(D)-Phe 1**). To test the **AND-Gate 1** probe, we incubated it with recombinant human CatL, B, S, or Casp3 either separately or sequentially in combinations (**Supplementary Fig. 1a**,**b**). Importantly, we found that the probe only produced a fluorescent signal when both cathepsin and Casp3 were incubated sequentially. We also found that both control probes remained non-fluorescent after incubation with both proteases, regardless of order of addition, thus validating the overall concept of the AND-gate strategy (**Sup Fig. 1c**,**d**). We then tested the stability of the linker using total tissue lysates derived from homogenized 4T1 breast tumors (**Sup Fig. 1e**). We found that, as expected, the **AND-Gate 1** probe was activated and the **(D)-Asp 1** negative control remained non-fluorescent. However, the **(D)-Phe 1** probe showed robust activation, suggesting a lack of stability at the amide bond connection between the Z-FK substrate and the linker.

Given the instability at the glutamine α-acid, we synthesized a 2^nd^ generation probe **AND-Gate 2** and the respective negative controls **(D)-Phe 2** and **(D)-Asp 2**, in which the natural (L)-Glu was replaced by the unnatural (D)-Glu (**Fig. 2b**). All peptide precursors were synthesized using solid-phase peptide synthesis (SPPS) using standard Fmoc chemistry (see Supporting Information for more details). The remainder of the synthesis was carried out in solution phase. The Z-FK substrate **1** was coupled to Fmoc-(D)-Glu(OAll)-OH using HCTU in the presence of 2,4,6-collidene in DMF. The allyl protected γ-carboxylic acid was subsequently removed using activated palladium in the presence of SiPhH_3_ to produce intermediate **2** with an overall yield of 76% over two steps. Next, **2** was coupled to the Casp3 substrate peptide **3** using the same coupling conditions mentioned above. After the coupling reaction was complete, piperidine was added to deprotect the Fmoc protected α-amine in one pot. The α-amine was then coupled to sulfo-Cy5-Osu, purified, and then deprotected in 80:20 TFA:DCM. The resulting deprotection afforded **4** in a 23% yield over the three step sequence. The two deprotected lysine side chains of **4** were coupled to sulfo-QSY21-OSu in the presence of excess DIPEA in DMSO over 48 h at 40 °C to produce **AND-Gate 2** in a 34% yield. The overall solution phase synthesis was six steps with an overall yield of 6%. The two negative controls **(D)Asp 2** and **(D)Phe 2** were synthesized in the same manner with similar overall yields.

We evaluated this 2^nd^ generation set of probes using the *in vitro* fluorogenic assay with recombinant proteases and found that the probes performed similarly to the first generation probes with only **AND-Gate 2** producing signal, and only after addition of both CatL and Casp3 (**Fig. 3a**,**b**). Using LC-MS analysis, we were able to detect the expected cleavage products after incubation with CatL or Casp3 alone, and with both proteases successively (**Supplementary Fig. 2**). Importantly, the second-generation probes containing the D-Glu linker showed good stability in tumor lysates, with both negative probes remaining inactive even after 2 hours of incubation (**Fig. 3c**). The **AND-gate 2** was also selective for Casp3 over other related initiator and executioner caspases (**Supplementary Fig. 3**).

**Figure 3.**
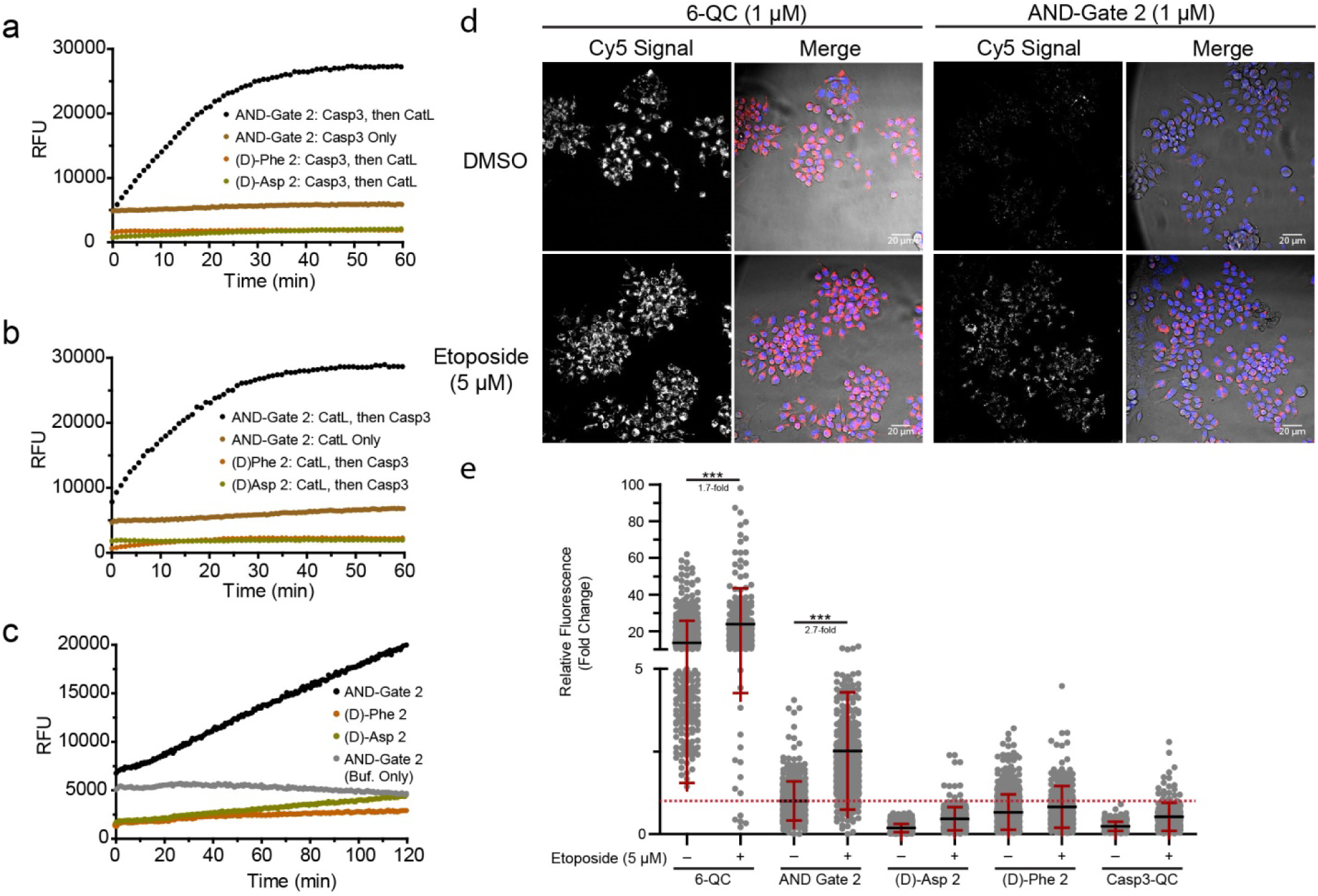
In vitro validation of the AND-gate strategy. (**a**) Progress curve of **AND-Gate 2** and negative control probes **(D)-Phe-2** and **(D)-Asp-2** after incubation with Casp3 (10 nM) alone or with Casp3 and then CatL (10 nM). All substrates were used at 10 μM. (**b**) Progress curves as in (**a**) except CatL was added first followed by cas3 (**c**) Progress curve over 2 hours incubation of the **AND-Gate 2** probe and corresponding negative controls, **(D)-Phe-2** and **(D)-Asp-2**, in either buffer or tumor lysate generated from 4T1 breast tumors in female BALB/c mice. (**d**) Representative fluorescent microscopy images of 4T1 cells co-cultured with RAW macrophages (1:1 ratio) labeled with either the single-parameter cathepsin probe **6-QC** or the **AND-gate 2** probe. Cells were incubated with either DMSO or etoposide (5 μM) for 24 h prior to addition of probes (1 μM). After 2 h, Hoechst stain was added to visualize nuclei and the cells were imaged. The Cy5 signal is shown in gray scale in the left panels and for the merged images on the right the brightfield is in grayscale. Blue is nuclear staining and red is the Cy5 probe signal. Representative images of cells incubated with control probes can be found in the Supplementary Information. (**e**) Scatter plot of fluorescent signal from (**d**) shown as the fold change of corrected total cellular fluorescence normalized to cells incubated with DMSO and **AND-Gate 2**. (Mann-Whitney Test, two-tailed *** p < 0.0001). Representative images of the negative controls **(D)Phe 2, (D)Asp 2**, and **Casp3-QC** can be found in the Supplementary Information.

Having solved the issue of linker stability, we tested the specificity of activation of the **AND-Gate 2** probe using a co-culture of breast tumor 4T1 cells and RAW macrophages. Cells were first incubated with DMSO or the cytotoxic agent etoposide (5 μM) for 24 h to induce apoptosis. The **6-QC, AND-Gate 2, Casp3-QC** (single substrate probe for Casp3, **Supplementary Fig. 4**), and respective negative control probes were added to the cells for 2 h before acquiring images (**Fig. 2d**). As expected, none of the negative control probes were activated under any conditions while **6-QC** was activated independently of apoptosis induction and **AND-Gate 2** was activated only in the apoptotic cells (**Fig. 3e**). These results confirm that the AND-Gate 2 probe required processing by both proteases to produce a fluorescent product that accumulates inside macrophage lysosomes.

**Figure 4.**
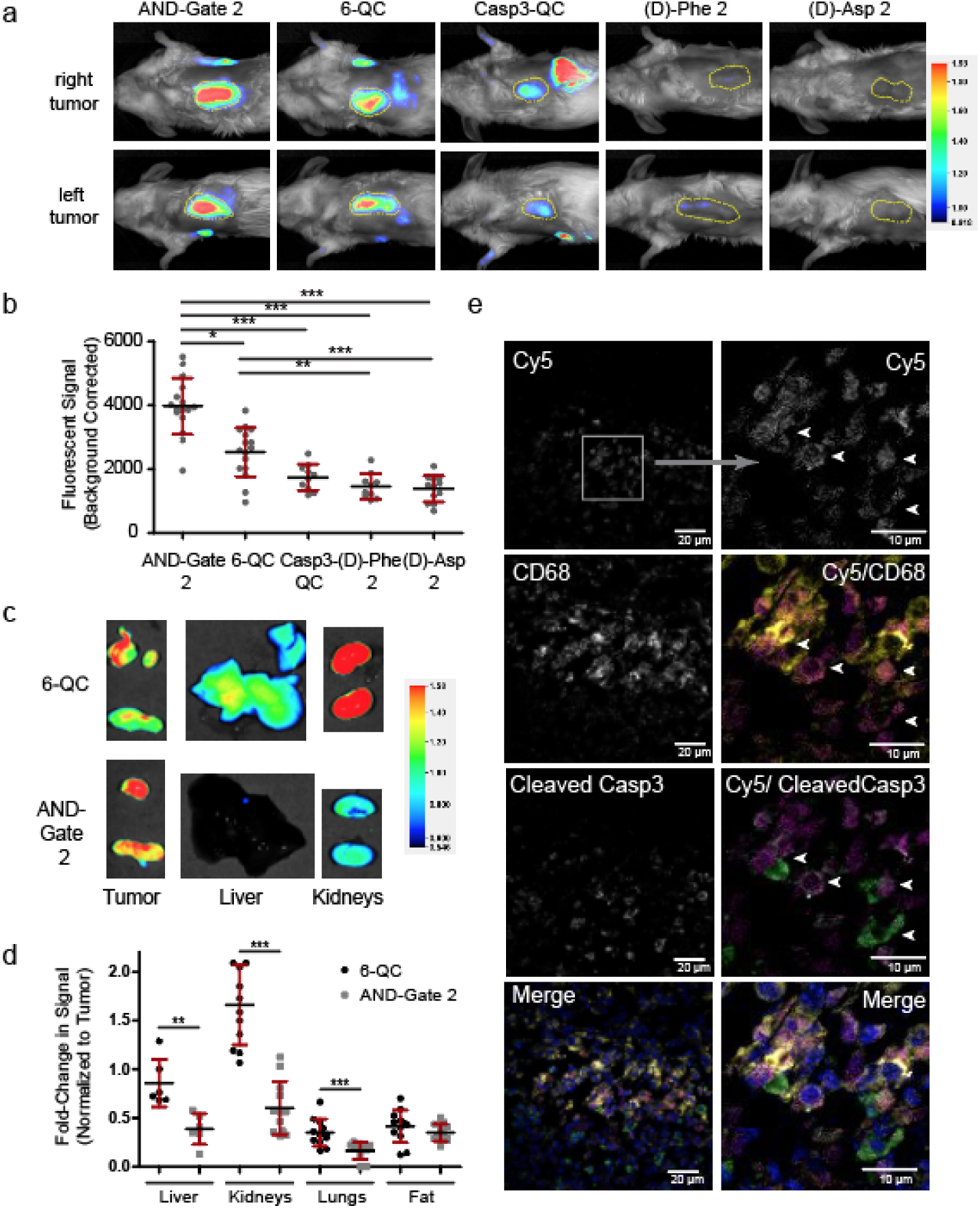
Enhanced selectivity and sensitivity of the AND-gate probe. (**a**) Images of BALB/c mice bearing 4T1 breast tumors 2 hr after probe injection (20 nmol). Fluorescent signal in tumors is displayed as rainbow plots and is normalized across representative images from each cohort. Tumor outlines are indicated with yellow dotted lines. (**b**) Scatter plot of average background corrected signal from all mice injected with each probe. (Tukey’s Test: *p < 0.05, **p < 0.01, ***p < 0.001). (**c**) Images of excised organs of mice after *in vivo* imaging (2 h post injection). Fluorescent signal is displayed as rainbow plots and is normalized between images. Images are representative of each cohort. (**d**) Scatter plot of the fold change in fluorescence signal compared to tumor signals for each tissue type in mice treated with the indicated probes. (Student’s T-test, two-tailed, Benforroni-Holm Procedure: **p < 0.01, ***p < 0.001). (**e**) Fluorescence microscopy images of 4T1 tumor sections from mice treated with the **AND-gate 2 probe** and stained for cleaved Casp3 and CD68 using specific antibodies. Single channel images are in grayscale. Merge images are as follows: Cy5 (probe) is magenta, CD68 is yellow, cleaved Casp3 is green, DAPI is blue. The same contrast and brightness settings were used to process each channel. Images of all live mice, organs, and additional examples of stained 4T1 tumor sections can be found in the Supplemental Information

### AND-Gate Probes have improved tumor selectivity in a mouse model of breast cancer

We next evaluated the **AND-Gate 2** probe and its respective negative controls in a 4T1 mouse model of breast cancer. We chose this model because it allows direct imaging of live animals using a NIR imaging system. The tumor-specific signal can be measured relative to surrounding background tissues and tissues can be subsequently removed for quantification and determination of overall probe activation in major organs distant to the tumor site. We imaged live animals 2 h post injection of probes and found that the **AND-Gate 2** probe showed strong tumor accumulation that resulted in a brighter tumor signal than the cathepsin only substrate **6-QC** (**Fig. 4a**). Importantly, the negative control probes produce no visible signals confirming the linker stability observed in vitro. Quantification of total tumor fluorescence confirmed that both the **AND-Gate 2** and single substrate probe **6-QC** produced a significantly increased signal compared to the negative controls **(D)-Phe 2** and **(D)-Asp 2**, and that this level of contrast over controls was several-fold higher for the AND-gate probe (**Fig. 4b**). The Casp3-specific substrate probe **Casp3-QC** that has only the caspase half of the AND-gate substrate showed a low-level tumor signal, however it was not significantly higher than the two negative control probes. The tumor-to-background ratio (TBR) for the **AND-Gate 2** was also significantly increased over the single-parameter **6-QC** probe (3.0 ± 0.3 compared to 2.3 ± 0.5; **Supplementary Fig. 5**).

**Figure 5.**
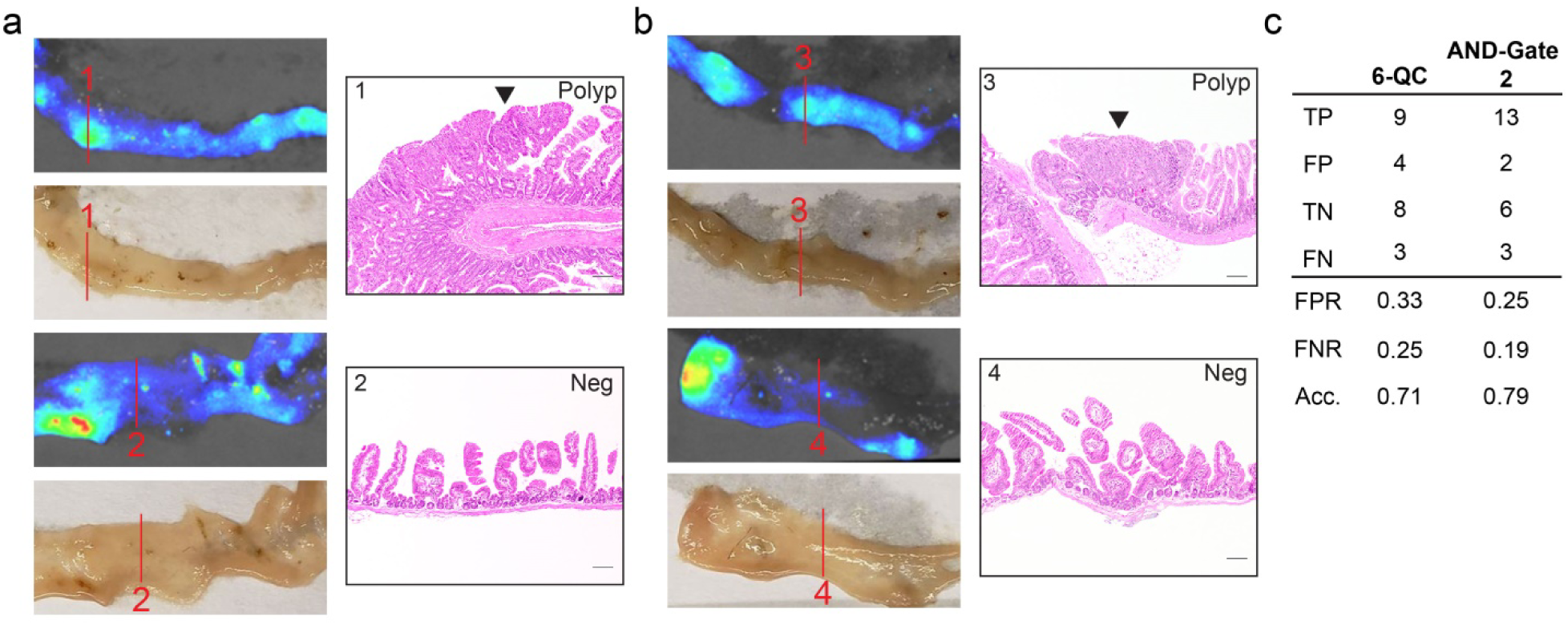
Imaging with the AND-Gate probe in a mouse model of colorectal cancer. Mice containing the Apc^min/+^ allele were aged 22 weeks and injected with **6-QC** or **AND-Gate 2** (R.O., 20 nmol). The mice were sacrificed 16 h post injection and tissues were linearized for fluorescent imaging. (**a-b**) Representative rainbow plot of fluorescent intensity, white light image, and H&E images of intestinal tissue from mice injected with (**a**) **6-QC** or (**b**) **AND-Gate 2**. Tissue was sectioned at numbered positions, H&E stained, and evaluated for the presence of tumor (sections 1 and 3 are examples of true positives; 2 & 4 are examples of true negatives). Arrowheads indicate polyps and scale bars = 100 μm. All images were contrast adjusted to show the difference between signal within polyps and normal tissue. (**c**) Areas in intestines with high, medium, or low fluorescence intensity were sectioned, H&E stained, and evaluated for the presence or absence of tumors. Sections were binned into four categories: true positive (contained tumor as expected, TP), false positive (unexpected tumor, FP), true negative (expected healthy tissue, TN), and false negative (unexpected healthy tissue, FN). Based on the sectioning results the false-positive rate (FPR), false-negative rate (FNR), and accuracy for each probe was calculated (n = 3 mice per probe). Rainbow plots of all data used for analysis, can be found in the Supplemental Information (**Supplementary Figure 9 & 10**).

In order to determine the overall tumor selectivity of the probes, we excised relevant organs for *ex vivo* imaging and quantified the background corrected signals. We found that the fluorescent signals in excised liver, kidneys, and lung tissue was significantly reduced in the animals treated with the **AND-Gate 2** compared to those treated with **6-QC** (**Fig. 4c**). Quantification of the fold change in fluorescent signals over tumor signals for each tissue confirmed the dramatic increase in selectivity of the AND-gate probe compared to the cathepsin-only **6-QC** probe (**Fig. 4d**). As a further analysis of probe activation *in vivo*, we fixed and embedded excised tumors for cryo-sectioning and histological analysis using fluorescence microscopy (**Fig. 4e**). We stained tumor sections for CD68, a marker for macrophages, as well as for cleaved Casp3. This analysis confirmed the presences of macrophages within the 4T1 breast tumors, as has been previously described^20^. We also observed numerous regions within the tumor tissues where cells contained active Casp3. These regions of apoptotic cells were always found directly adjacent to, or co-localized with, macrophage cell populations. The brightest probe signal was always found at the intersection between macrophages and cells with active Casp3 confirming that Casp3 and Cats are both active within a tumor and generally found in close proximity making them ideal targets for use in the AND-gate strategy.

### AND-gate probes have improved false positive and false negative rates in a mouse model of colorectal cancer

To further demonstrate the general applicability of the AND-gate strategy to other tumor models, we compared the performance of **AND-Gate 2** to the single substrate cathepsin probe **6-QC** in an Apc^min/+^ colorectal cancer mouse model. This model provides an opportunity to test the AND-Gate probe in a spontaneous cancer model containing a genetically distinct cancer from that of the 4T1 breast tumor model. Polyps in this model are difficult to distinguish from healthy tissue by eye due to their small size. Mice containing polyps were injected with each probe and sacrificed 16 h post injection. Overall, both probes were able to fluorescently label polyps in mice after systemic injection (**Fig. 5**). Using the fluorescent signal from the probes, we chose areas of high, medium, and low fluorescent signal to be sectioned, stained for hematoxylin and eosin (H&E), and evaluated for the presence of tumor. Areas containing high or medium fluorescence intensity were expected to contain tumor while areas with low fluorescence intensity were expected to be healthy. The evaluated sections were determined to be true positive (TP), false positive (FP), true negative (TN), or false negative (FN) based on the expected and actual outcomes for the presence of tumor (**Fig. 5c**). This analysis determined that the **AND-Gate 2** probe had a lower false positive rate (FPR) and false negative rate (FNR) compared to **6-QC** (0.25 vs 0.33 FPR and 0.19 vs 0.25 FNR). Based on this analysis, the two probes are comparable in their ability to highlight polyps in this colorectal cancer mouse model, but the **AND-Gate 2** has a higher accuracy compared to the single substrate probe **6-QC**.

### Synthesis of 780 nm emitting AND-Gate probe and evaluation using the da Vinci^®^ Xi Surgical System

To evaluate the performance of our AND-Gate probe strategy during robotic fluorescence-guided surgery, we synthesized an AND-Gate probe containing a fluorophore that has excitation and emission wavelengths compatible with the FDA approved Firefly^®^ detection system on the da Vinci^®^ Xi Surgical System. Currently, the Firefly^®^ fluorescence detection system is specifically tuned to the excitation/emission properties of indocyanine green (ICG, ex/em: 780/820 nm). We initially sought to synthesize an AND-Gate probe containing the conjugatable version of ICG (containing a carboxylic acid) and the respective quencher QC-1, which is non-fluorescent and has strong absorbance between 600-900 nm. However, conjugating ICG to the peptide scaffold caused unexpected reactivity issues possibly caused by aggregation or tertiary folding of the molecule. Therefore, we used a heptamethine cyanine fluorophore FNIR-Tag that was designed by the Schnermann group to resist aggregation and have better water solubility compared to ICG^32^. We conjugated the FNIR dye successfully to the α-amine on the central (D)-Glu linker and subsequently attach two QC-1 quenchers to the lysine side chains to produce **AND-Gate-FNIR** (**Supplementary Scheme 2**). Although this fluorophore has slightly lower excitation/emission maximum wavelengths compared to ICG (765/788 nm), the fluorescence can be detected by the Firefly^®^ detection system, albeit with lower sensitivity than for ICG. We confirmed that **AND-Gate-FNIR** was efficiently processed by CatL and Casp3 using the recombinant proteases in a fluorogenic substrate assay (**Supplementary Fig. 12**). Like the Cy5 AND-Gate probes, **AND-Gate-FNIR** produced a fluorescent signal only after incubation with both proteases, regardless of order of addition.

We then compared the performance of **AND-Gate-FNIR** to the single substrate probes **6-QC-ICG** and **6-QC-NIR** during robotic surgery using mice bearing 4T1 breast tumors. Both single substrate probes were developed in our laboratory and contain either an ICG or Dylight780-B1 (NIR) fluorophore, which are activated by cysteine cathepsin proteases^25^. We injected mice bearing 4T1 breast tumors with probes 2 h prior to performing robotic surgery (20 nmol, I.V.). The **AND-Gate-FNIR** produced fluorescent contrast in the tumor compared to the surrounding tissue, which clearly highlighted the margin during surgery (**Fig. 6a**). Despite using a fluorophore with lower excitation/emission wavelengths compared to ICG, the **AND-Gate-FNIR** had comparable fluorescent signal intensity to the **6-QC-ICG** probe and much improved signal in comparison to **6-QC-NIR**, which also has lower excitation/emission wavelengths than ICG (788/799 nm).

**Figure 6.**
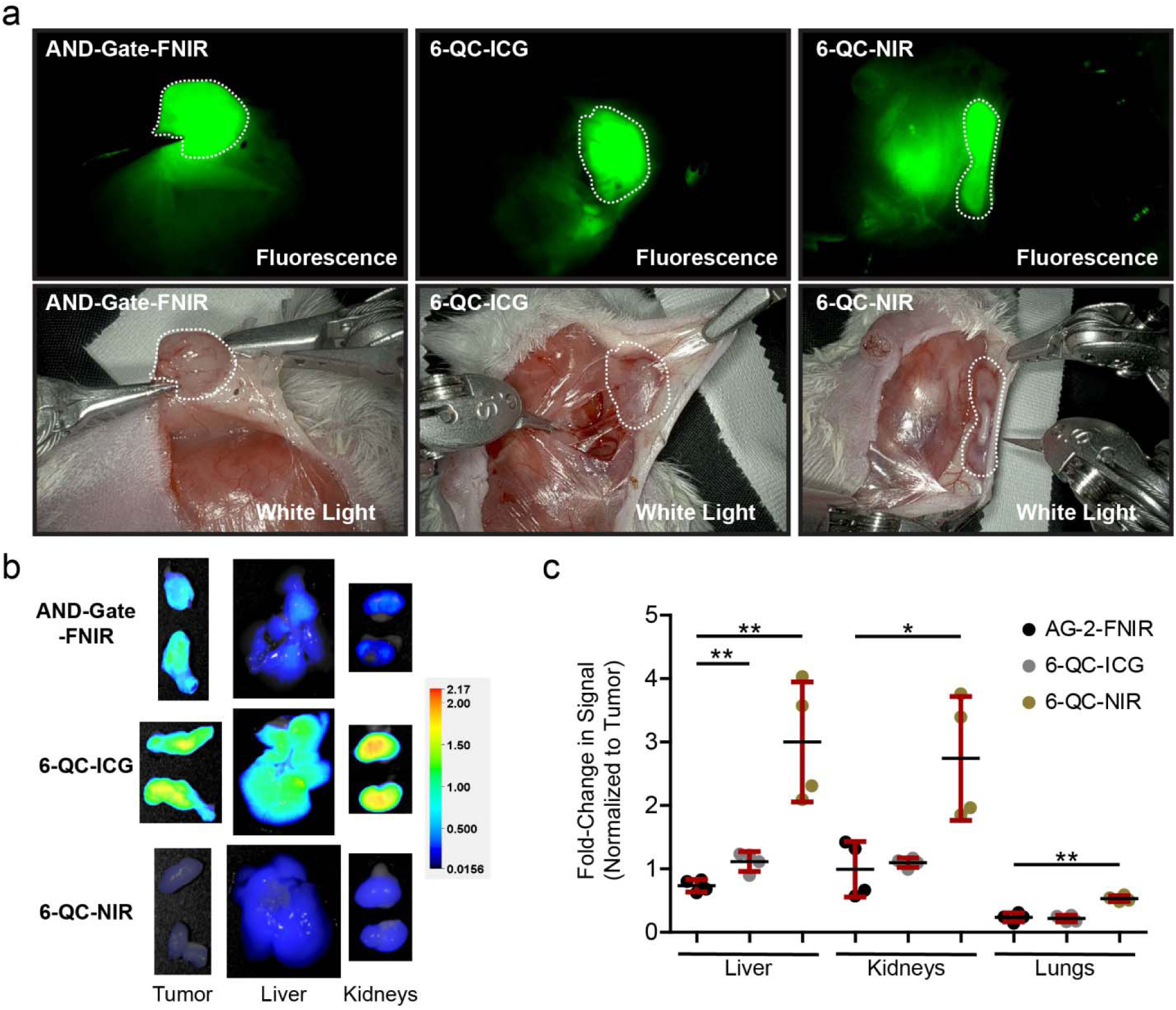
Images from robotic fluorescence-guided surgery with the da Vinci^®^ Xi Surgical System equipped with Firefly^®^ detection and quantification of fluorescence in healthy organs. (**a**) Images of 4T1 breast tumors in mice injected with **AND-Gate-FNIR, 6-QC-ICG**, or **6-QC-NIR** (20 nmol, I.V.). Tumors are outlined with white dotted line. (**b**) Representative images of excised tumors and healthy organs. Signals are normalized and displayed as rainbow plots. (**c**) Relative quantification of fluorescent signal in liver, kidneys, and lungs normalized to tumor signal (Student’s T-test analysis, two-tailed: *p < 0.05, **p < 0.01).

Following robotic surgery, tumors and organs from mice injected with each probe were imaged *ex vivo* using the LiCor Pearl imaging system to quantify the fluorescent signal. Representative images of excised tumors and organs confirmed a reduced signal in the liver and kidneys compared to the signal found in the tumors for **AND-Gate-FNIR** compared to both of the 6-QC probes (**Fig. 6b**). Quantification of the signal in the organs normalized to the tumor signal confirmed a significant reduction in background in the liver for mice injected with **AND-Gate-FNIR** compared to both 6-QC probes and a significant reduction in background for all healthy organs compared to **6-QC-NIR** (**Fig. 6c**).

To further demonstrate the utility of the **AND-Gate-FNIR** probe, we performed a resection of a primary subcutaneous mammary tumor and subsequently used probe fluorescence to assess the resulting subcutaneous tumor bed. The residual cancer cells left after excision of the primary tumor were not visible by white light imaging but could easily be visualized using the Firefly^®^ detection of the probe signal (**Supplementary Movie 1; Fig. 7**). We then removed the subcutaneous tumor bed and overlying haired skin with significant surrounding probe-negative tissue. This tissue was formalin-fixed and sectioned such that H&E slides contained both probe positive and probe negative regions on a single slice. Analysis of the H&E stained tissues by a board-certified pathologist confirmed that the regions showing probe activation contain residual tumor left behind following excision, while probe negative regions contained normal subcutaneous and cutaneous tissue. Thus, the **AND-Gate-FNIR** probe was able to accurately detect areas of residual tumor cells left behind after excision of a bulk tumor.

**Figure 7.**
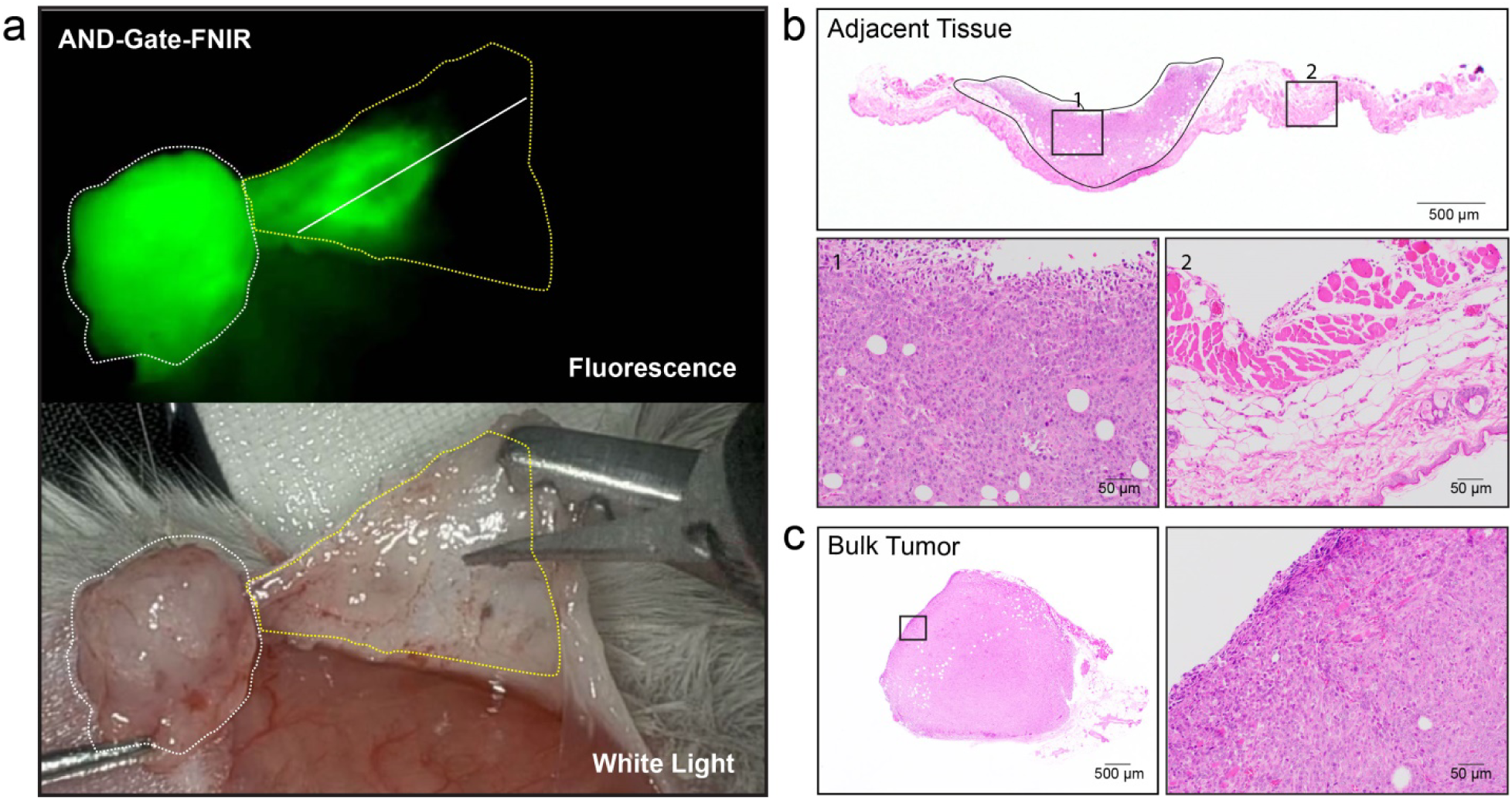
Detection of residual tumor margins using the **AND-Gate-FNIR** probe and Firefly® detection system (**a**) Screen capture images (Firefly® and white light) of a primary excised tumor (white dotted line) and the resulting adjacent tissue (yellow dotted line). The remaining tumor bed (including the subcutaneous and cutaneous tissue) was excised for sectioning. Histology was performed on sections of the tumor bed (solid white line indicating position of histology section) and the bulk tumor. Sectioning was performed perpendicular to the plane of the image. (**b**) H&E staining of adjacent probe-positive and probe-negative tissue showing 20x magnification images of regions of residual tumor (area 1) and surrounding healthy tissue (area 2). The black outline indicates area of tumor within the section. (**c**) H&E staining of a section taken from the bulk tumor with a 20x magnification of the region shown in the box area.

## Discussion

The sensitive and accurate detection of tumor margins is a central goal for effective treatment of any type of solid tumor using surgery. Currently, there are a number of effective ways to image the location of a tumor in the body. However, most of the current imaging strategies are not suitable for real-time applications during surgery. Furthermore, they often depend on contrast agents that lack a level of specificity that allows them to be used in diverse cancer tissue types. Recent efforts to address these limitations in current imaging technologies have focused on targeted contrast agents that produce fluorescent signals only within the tumor microenvironment. While this approach has proven valuable and several agents are currently in human clinical trials, there still remains a need to find new strategies to improve the selectivity of optical contrast agents so that they can be used in virtually any type of tumor resection. We and others have developed activatable ‘smart probes’ that turn on a fluorescent signal when processed by a protease that is highly activated in tumor tissues. These reagents provide overall high contrast in some tumor types where levels of the active protease in the surrounding normal tissues is low. Here, we outline a general strategy to generate a next-generation of optical smart probes that require sequential processing by multiple tumor-associated enzymes. This multivariate AND-gate type sensor has the potential for highly increased tumor selectivity by incorporating substrates that are specifically processed by enzymes that exist in normal tissues but that are only found together in the tumor microenvironment. Using this strategy, background signal in healthy surrounding tissue is eliminated, resulting in high image contrast and clear margin detection even for probes with overall low signal strength.

We chose to build the first AND-ate probe using two substrates. The first is a sequence that have previously shown to work well for multiple cysteine cathepsins and that is used in our single substrate tumor imaging probe **6-QC**^20,25^. For a second substrate, we chose to use a peptide sequence that has been optimized for cleavage by Casp3^33,34^. We chose these enzymes because of their overall distinct and non-overlapping primary sequence specificity. Furthermore, while all tissues contain some level of active cathepsins, Casp3 is only activated in the late stages of apoptosis, which typically does not take place in healthy tissues. Multiple types of cell death are activated within tumors as the result of several factors including nutrient and oxygen starvation. Dying cells within the tumor activate infiltrating immune cells, such as macrophages that then clear the cells by lysosomal engulfment. Therefore, there exists a specific niche within the tumor microenvironment where both lysosomal and apoptotic proteases co-exist, making these proteases ideal choices for use in tumor-specific AND-gate sensors.

Because we have designed a general strategy in which the fluorescent reporter is attached to a central linker that is the final end product of probe processing, it is possible to further engineer our first generation AND-gate probes to respond to additional enzymatic processes beyond a binary set of targets. This hub and spoke model should also be amenable to further optimization of chemistry to allow production of diverse AND-gate sensors by swapping of the spokes around the central fluorescent hub. In addition, it is possible to replace the current quencher/fluorophore strategy with a FRET fluorophore pair that would allow ratiometric imaging to detect probe distribution as well as activation. This strategy would help to reduce heterogeneity of imaging signals due to differential probe distribution while also reducing background signals by allowing quantification based on a ratio of signals, which can be a large value even when probe concentrations are low. We are currently working to develop ratiometric AND-gate probes as well as multivariate versions that respond to more than two proteases. There are a number of additional proteases that have been shown to be preferentially activated in solid tumors of diverse origins, and many of these proteases have defined substrate specificity that would make design of next generation AND-gate sensors straightforward.

In addition to using the multivariate AND-gate strategy to improve imaging contrast, it should in theory also be possible to apply this approach for selective drug delivery applications. Such a strategy would require attachment of an individual or multiple drugs to the central hub through linkages that block activity of the agents and that can be processed within a specific tissue location. One could envisage such an AND-gate therapy that releases one or more active drugs only when multiple proteases cleave the linkages tethering the molecules to one another. We are currently exploring approaches to use the AND-gate approach for such applications.

We show here that the current AND-Gate FNIR probe is compatible with the existing FireFly detection on the widely used da Vinci Surgical System. Our results suggest that these probes could be rapidly integrated into existing surgical workflows to enable visualization of residual tumor tissue remaining after removal of a primary mass. While the 4T1 breast cancer model produces tumors that are highly encapsulated making them easily visualized by standard white light imaging, we show here that after careful removal of the primary mass, a significant number of tumor tissue remains at the skin/tumor interface. Using standard histology methods, we show that the probe accurately highlights these regions of residual cancer cells, even when those areas contain a low number of cells that cannot be detected using white light imaging. These results suggest that the AND-gate strategy will enhance the effectiveness of tumor resections using the da Vinci Surgery System even in locations where high background signal would normally limit effective visualization of the margin.

## Conclusion

Overall, the studies presented here provide a roadmap for the design of optical contrast agents that respond to unique enzymatic signatures that are indicative of a disease state. In the current design, we demonstrate the utility of the AND-gate approach using substrates that are uniquely processed by two proteases, which are both activated in the tumor microenvironment. The design of the AND-gate protease probe used here required engineering of the linker group to prevent activation by undesired protease activities found in normal healthy tissues but ultimately resulted in a contrast agent that both improved tumor-selectivity and also increased tumor uptake resulting in a brighter overall signal. We are currently working to further optimize the central linker such that higher order probes can be rapidly generated by attachment of various substrates containing quenchers using orthogonal chemistry. This should allow synthesis of probes with high selectivity for diverse cancer types based on proteolytic signatures of those tumors. It also should, in principle, enable generation of probes that respond to other classes of enzymes that are capable of processing substrates to liberate a central fluorescent fragment.

## METHODS

### Chemistry Methods

See Supporting Information for synthetic protocols.

### Biology Methods

#### General Cell Culture

4T1 cells and RAW246.7 macrophages were cultured separately as previously describe^20,35^. RAW246.7 macrophages were cultured in Dulbecco’s Modified Eagle’s medium (DMEM) supplemented with 10% fetal bovine serum (FBS), and 100 units/ml penicillin and 100 μg/mL streptomycin. 4T1 cells were cultured in Roswell Park Memorial Institute (RPMI) 1640 medium with L-glutamine supplemented with 10% FBS, and 100 units/mL penicillin and 100 μg/mL streptomycin.

#### Fluorogenic Substrate Assay

All proteases were active site titrated as previously described to obtain active protease concentrations^20^. The buffers used for caspases and cathepsins were made as previously described^20,36^. The reducing agent 1,4-dithiothreitol (DTT) was freshly added to buffers just prior to use. All assays were conducted in black, opaque flat bottom 384-well plates (Greiner Bio-One). For single substrate probes and first cleavage of AND-Gate probes, compounds were diluted in the respective protease buffer from 10 mM stock solutions in DMSO, and 15 μL of substrate (20 μM) was added to each well. Immediately prior to beginning fluorescent measurements, 15 μL of protease was added to wells containing substrate using a multichannel pipette. All proteases were at a final concentration of 10 nM in the well unless otherwise indicated. Fluorescent measurements for probes containing a sulfo-Cy5 were read above the well with a Biotek Cytation3 Imaging Reader (7.00 mm read height, gain = 100, ex/em = 640/670 nm, normal read speed). Fluorescent measurements for probes containing ICG or FNIR-Tag fluorophores were read above the well with a SpectraM2 plate reader (7.00 mm read height, ICG: ex/em 780/820 nm, FNIR-Tag: ex/em 765/790 nm, normal read speed).

After confirming AND-Gate probes are not activated upon single addition of protease, probes were incubated with single proteases in an Eppendorf tube prior to second protease addition and fluorescent measurements. AND-Gate probes or negative controls (40 μM) were initially digested in 20 nM protease in the respective protease buffer. After 2 h at 37 °C, reactions with cathepsin buffer (pH 5.5) were exhausted by addition of 2 M NaOH (1.4 μL/100 μL of buffer) and then diluted with caspase buffer to obtain a concentration of 20 μM total substrate (pH 7.0). Reactions containing caspase buffer (pH 7.0) were exhausted by addition of 1 M HCl (2 μL/100 μL of buffer) followed by dilution with cathepsin buffer to obtain a final concentration of 20 μM total substrate (pH 5.5). The singularly processed probes were added to wells (15 μL) and the second protease was added just prior to beginning fluorescent measurements (15 μL, 10 nM final protease concentration, 10 μM final substrate concentration). Fluorescent signal over time was measured as mentioned above.

#### Live Cell Fluorescent Microscopy Assay

RAW246.7 macrophages and 4T1 cells were diluted to 1 x 10^5^ cells/mL with their respective media and then mixed in a 1:1 ratio. The mixture of cells were seeded in a 96-well, half-area, μclear® bottom, cell culture plate (Greiner Bio-One) at 30 μL per well, and allowed to adhere to the bottom of the plate for 24 h. Next, cells were incubated with DMSO (1 % v/v) or etoposide (5 μM). After 24h with DMSO or etoposide, probes were added to the cell media (5 μL, 1 μM) and incubated for 2 h. Then, Hoescht 33342 was added to the media (5 μg/mL) and the cells were imaged using a 40X oil emersion objective on a Zeiss Axiovert 200 M confocal microscope. Acquired Z-stacks consisted of 24 16-bit images taken in 1 μm increments apart beginning and ending 12 μm from the focal plane. The channels used were set on separate tracks. The channel settings were as follows: Cy5-Gain = 545, laser ex = 639 nm, 2% power, Hoescht 33342-gain = 750, laser ex = 405 nm, 4% power, and DIC-(T-PMT from Cy5 channel, gain = 480.

Images were processed using ImageJ. Images shown are flattened from Z-stacks by taking the maximum signal in each pixel. All images are adjusted to the same contrast and brightness levels. The corrected total cellular fluorescence (CTCF) was determined by manually tracing cells based on the DIC and nuclear staining images, and measuring the total fluorescence within each cell from the flattened Z-stack images. CTCF was calculated according to the following formula: CTCF = Integrated density – (area of cell x average background). The average background was obtained by tracing circles where no cells were present and averaging the overall signal. Cells that were overlapping or on the edge of the image were not counted. Sample size (n) for each condition: 6-QC, DMSO: 416; 6-QC, 5 μM Etoposide: 266; AND-Gate 2, DMSO: 399; AND-Gate 2, 5 μM Etoposide: 462; (D)-Phe 2, DMSO: 402; (D)-Phe 2, 5 μM Etoposide: 269; (D)-Asp 2, DMSO: 599; (D)-Asp 2, 5 μM Etoposide: 246; Casp3-QC, DMSO: 226; Casp3-QC, 5 μM Etoposide: 338. CTCF measurements were normalized to **AND-Gate 2**, DMSO treated control to obtain fold-change values.

#### Animal Models and Fluorescent Imaging

All animal care and experimentation was conducted in accordance with current National Institutes of Health and Stanford University Institutional Animal Care and Use Committee guidelines. All *in vivo* and *ex vivo* imaging was conducted with a LiCor Pearl Trilogy imaging system using the 700 nm and white light image settings with a resolution of 170 μm. Live mice were imaged at indicated time points after R.O. or I.V. tail vein injection of probe under isoflurane anesthesia. After imaging, mice were sacrificed under isoflurane anesthesia by cervical dislocation. Each measurement represents an independent, distinct sample.

#### 4T1 Breast Tumor Model

4T1 cells (ATCC) suspended in 1X PBS (100 μL, 1 x 10^6^ cells/mL) were injected subcutaneously into the 3 and 8 mammary fat pads of 6-8 week old BALB/c ByJ female mice (Jackson Laboratory). Mice were monitored for tumor formation and were injected with probe for imaging between 7-10 days after seeding 4T1 cells. Probes were dissolved in 1X PBS (10% DMSO) and injected I.V. tail vein (100 μL, 20 nmol). Hair was removed to expose the tumors with Nair® lotion while the mice were under anesthesia 1 h prior to imaging. Mice per probe used for imaging analysis: AND-Gate 2: 5; 6-QC: 8; (D)-Phe 2: 5; (D)-Asp 2: 7; Casp3-QC: 5. Data was collected from three technical replicates where each probe was injected into at least one mouse for each replicate. Images were processed using LiCor Imaging Studio (Ver 5.2). Background corrected signal was obtained by drawing circles around tumors and then subtracting the signal directly adjacent to the tumor without hair. The areas of circles used to calculate the background corrected signal was the same for each tumor. Tumor-to-background (TBR) was calculated by placing two circles of the same size over the tumor and the exposed skin directly adjacent to the tumor and dividing the two fluorescent measurements.

#### Histological Analysis of 4T1 Tumor Sections using Confocal Fluorescence Microscopy

4T1 breast tumors in the mammary fat pads of BALB/c mice injected with probe were excised postmortem. The tumor tissues were fixed in neutral buffered formalin solution (4% formaldehyde) for 24 h at 4 °C. The fixed tumor tissues were then soaked in 30% w/v sucrose in 1X PBS for 24 h at 4 °C. Tumor samples were embedded in Optimal Cutting Temperature (O.C.T.) compound and kept at -80 °C until further use. Embedded tissues were then crysosectioned (5 μm) onto glass slides. The samples were stored at -20 °C prior to staining. The cryosections were first washed with 1X PBS 3X in a slide chamber. Then, the samples were blocked with 3% w/v BSA for 24 h at 4 °C. Samples were washed 3X with 0.5% w/v BSA, then tissues were outlined with a hydrophobic pen, followed by incubation with 1:2,500 dilution of primary antibodies for CD68 (rat anti-mouse, ABD SeroTec) and cleaved Casp3 (rabbit anti-mouse, Cell Signaling) in 0.5% w/v BSA in 1X PBS for 24 h at 4 °C. Samples were then washed 5X with 0.5% w/v BSA in 1X PBS. Tissues were then incubated with anti-rat alexafluor488 and anti-rabbit alexafluor594 secondary antibodies (Invitrogen) diluted 1:5,000 for 1 h at RT. Tissues were washed 5X with 0.5% w/v BSA in 1X PBS, followed by 3X washes with 1X PBS. Tissues were mounted with Vectashield mounting medium containing DAPI.

Sections of tumors were imaged using a 40x oil emersion objective on a Zeiss Axiovert 200 M confocal microscope. Images are of a single focal plane (16-bit). All channels were set on separate tracks for imaging. The channel settings were as follows: Cy5–Gain = 730, laser ex = 639 nm, 20% power; A594(cleaved Casp3)–gain = 600, laser ex = 555 nm, 4% power; A488(CD68)–gain = 640, laser ex = 488, 4% power; DAPI–gain = 600, laser ex = 405 nm, 2% power.

#### Apc^min/+^ Colorectal Cancer Model

Apc^min/+^ on the C57BL/6 background were purchased from Jackson Laboratories and maintained in a specific pathogen-free facility. Mice were allowed to age to 22 weeks to allow for polyps to develop before the experiment. Mice were injected with **6-QC** or **AND-Gate 2** (R.O., 20 nmol, 100 μL) in 1X PBS containing 10 v/v % DMSO. Mice were euthanized 16 h post injection. Following euthanasia, small intestinal tissues were excised from the level of the stomach to the cecum. The intestine was linearized parallel to its longitudinal axis for visualization of mucosal polyps and placed on an index card (serosal surface down) to retain linear orientation. Brightfield and fluorescent imaging was performed as described above using a LiCor Pearl Imaging system. Following imaging, tissues were immersion fixed in 10% neutral buffered formalin for 72 hours at room temperature and transferred to 70% ethanol.

#### Histopathological Analysis of Apc^min/+^ mouse colons

Intestinal tissues were grossly compared to fluorescent images for identification of targeted lesions. Areas of high, medium, and low fluorescent signal were chosen for sectioning and analysis. It was expected that areas containing high and medium fluorescent signal intensity would be positive for tumor tissue and areas of low signal would be healthy tissue. In total, n=50 distinct locations were selected (n=25 sections per probe) for histopathologic evaluation based on fluorescent signaling. Tissue sections were cut perpendicular to the longitudinal axis of the intestine, processed routinely, embedded in paraffin, sectioned at 5 μm, and stained with hematoxylin and eosin (H&E). All sections were blindly evaluated by a board-certified veterinary pathologist for the presence or absence of mucosal polyps. If histopathologic results deviated from expected results based on fluorescent imaging (n=10), paraffin blocks were melted, rotated 90°, re-imbedded, and H&E stained to evaluate for polyps in various planes of section.

#### Robotic Fluorescence-Guided Surgery

Breast tumor-bearing mice (4T1) were administered the indicated probes via I.V. tail vein (100 μL, 20 nmol in 30% v/v PEG400, 10% v/v DMSO in 1X PBS) 2 h prior to surgery. Robotic fluorescence-guided surgery was performed with an FDA approved da Vinci® Xi Surgical System equipped with a Firefly® fluorescence detector. Surgery was initially performed under inhaled isoflurane and injection of a mixture of ketamine/telazol solution. After resection of the tumors, mice were sacrificed by injection of ketamine under heavy anesthesia, followed by cervical dislocation. Breast cancers were detected with a combination of white light and fluorescence signal based on probe activation as a guide to determine tumor margins from healthy tissue. Videos of the surgical procedures are available in the Supporting Information. For comparison of fluorescent signal in *ex vivo* tumors and organs: n = 3 mice per probe.

#### Histopathological analysis of 4T1 Tumors

Tissues including the bulk 4T1 breast tumor and tissue adjacent to the tumor were excised, fixed in 4% formaldehyde in buffered solution, processed routinely, sectioned at 5 μm, and stained with H&E. Adjacent subcutaneous and cutaneous tissues were embedded in paraffin with the subcutaneous side down (bulk tumor adjacent), such that each slide contained both probe-positive and probe-negative regions. Step sections were taken through the entirety of the tissue at 100 μm intervals.

#### Data Availability

All data and information necessary to reproduce the results reported in this manuscript are provided. Any additional data that support the findings of this study is available upon reasonable request.

## Supporting information

Widen et. al. Supplemental Information

## ACKNOWLEDGEMENTS

This work was supported by NIH grants R01 EB026285 (to M.B.) and a Stanford Cancer Institute Translational Oncology Program seed grant (to M.B.), American Cancer Society–Grand View League Research Funding Initiative Postdoctoral Fellowship, PF-19-105-01-CCE (To J.W.), DFG Research Fellowship TH2139/1-1 (to M.T.), and Stanford ChEM-H Chemistry/Biology Interface Predoctoral Training Program and NSF Graduate Research Fellowship Grant DGE-114747 (to J.J.Y). Special thanks to Scott Snipas in the Guy Salvesen Lab at Sanford Burnham Prebys Medical Discovery Institute for gifting the recombinant caspases used in this study. Thanks to the Turk Lab at J. Stefan Institute for providing the recombinant cathepsin proteases used in this study. Thanks to Stacy A. Malaker and Nick Riley in the Carolyn Bertozzi Lab at Stanford University for the high-resolution mass analysis of the AND-Gate probes. Thanks to Michael P. Luciano and Martin J. Schnermann at the National Cancer Institute for supplying the FNIR-Tag-OSu used to synthesize the **AND-Gate-FNIR** probe. Thanks to the Peter Santa Maria Lab for using their SpectraM2 plate reader.

## AUTHOR CONTRIBUTIONS

M.B. and J.C.W conceived of the AND-Gate probe concept and designed all experiments. J.C.W synthesized all AND-Gate probes, conducted the fluorogenic substrate assays, live and fixed cell fluorescent microscopy experiments, and mouse model experiments. J.C.W and M.B. wrote the text of the paper and constructed the figures with input from J.J.Y. and K.C.M. M.T. and J.J.Y helped perform live and *ex vivo* imaging during the 4T1 cancer mouse model experiment including dissection of the mice. M.T. aided in the immunohistochemical analysis of 4T1 tumors. A.A, A.K., and J.S. assisted with the robotic surgery. S.R. assisted in the colorectal cancer mouse model. K.M.C. evaluated H&E sections for the colorectal and 4T1 breast cancer mouse models.

