## Supplementary material for "Multivariate AND-gate substrate probes as enhanced contrast agents for fluorescence-guided surgery": Widen et. al. Supplemental Information

TABLE OF CONTENTS FOR SUPPLEMENTARY DATA

***Page***


I. **Supplementary Schemes** ........................................................................ S3

II. **Supplementary Figures** ........................................................................ S5

III. **Chemistry Methods**.............................................................................. S15

IV. **HPLC Purity Analysis** .......................................................................... S22

V. **Supplementary References**.................................................................. S26

### I. Supplemental Schemes

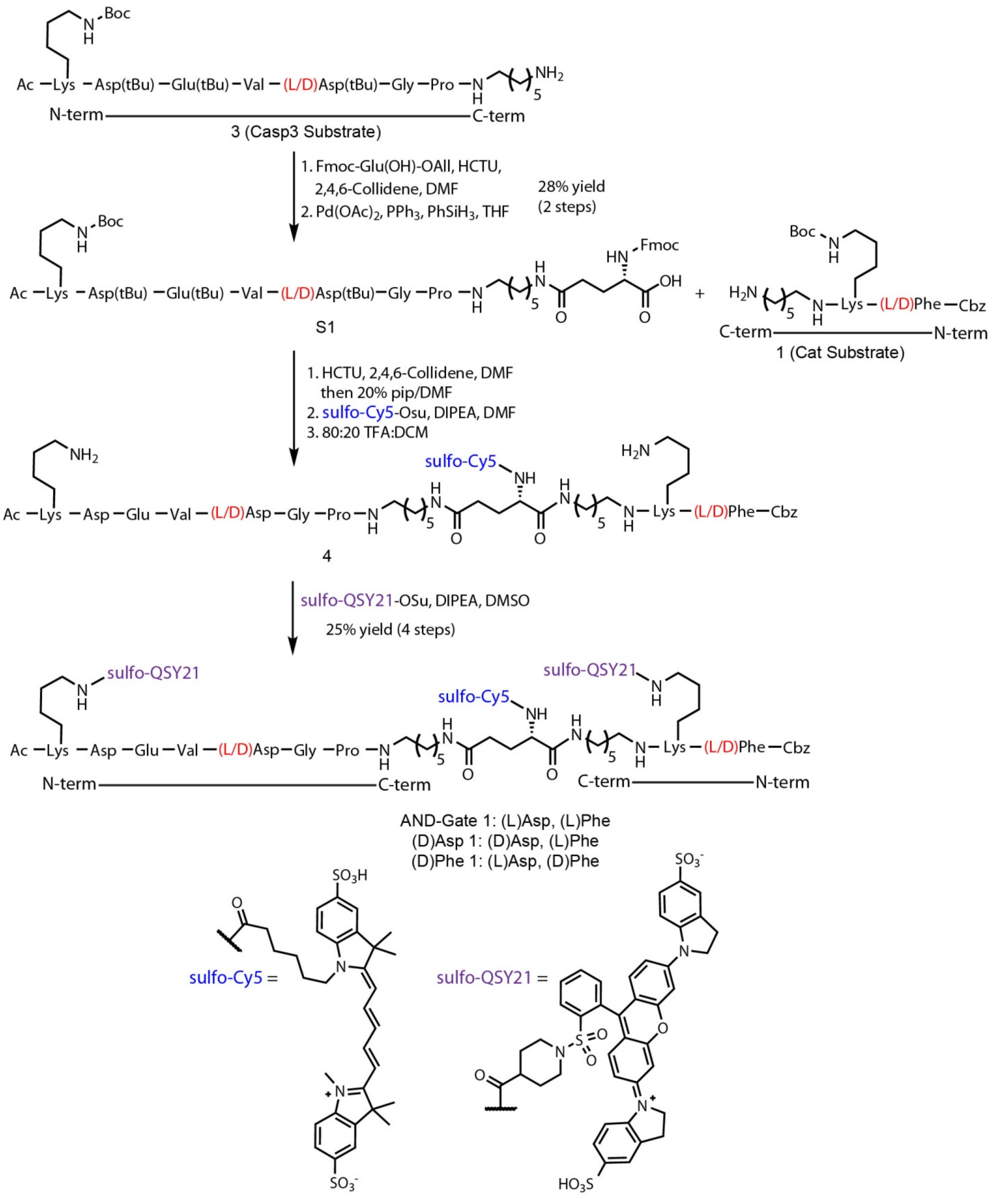


**Supplementary Scheme 1.** Synthesis of **AND-Gate 1**, **(D)Asp 1**, and **(D)Phe 1** containing an (L)-Glu central linker. See Methods section for synthetic details.

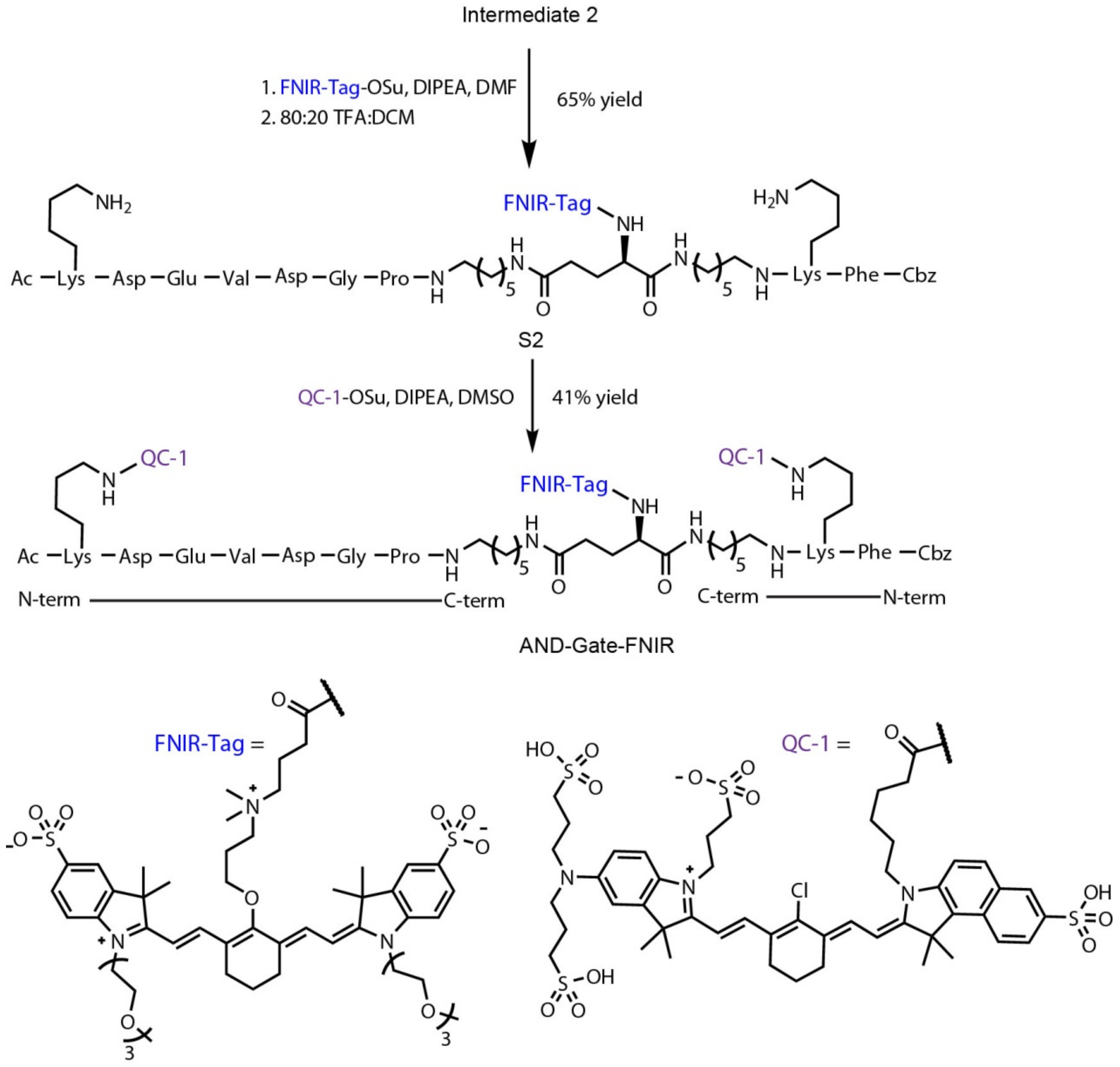


**Supplementary Scheme 2.** Synthesis of **AND-Gate-FNIR** containing a (D)-Glu central linker with an FNIR-Tag[1] fluorophore and QC-1 quencher system. See Methods section for synthetic details.

### II. Supplemental Figures

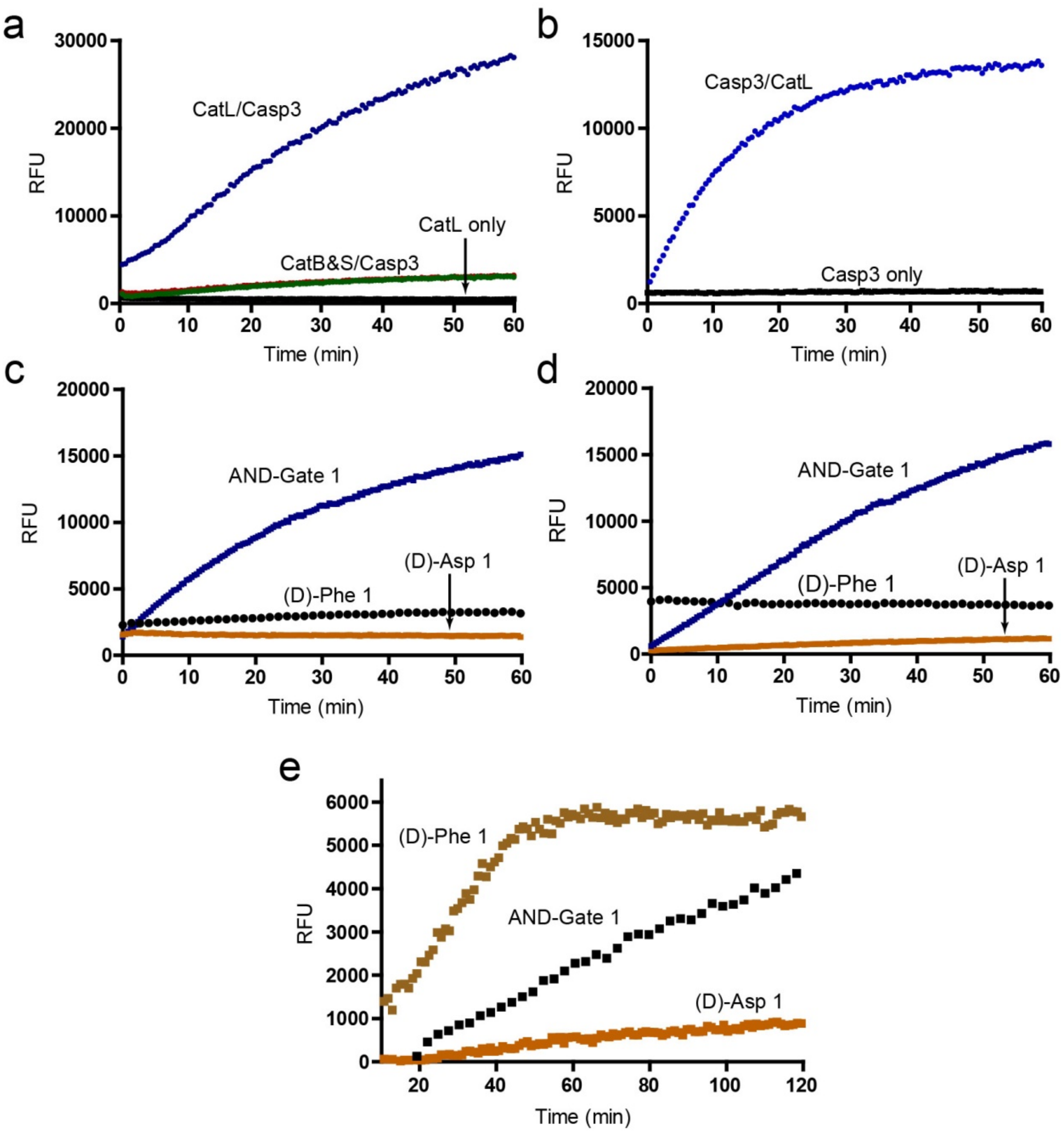


**Supplementary Figure 1.** Evaluation of **AND-Gate 1** and respective negative controls with human recombinant proteases in a fluorogenic assay. (**a**) **AND-Gate 1** was first incubated with Cat L, B, or S (10 nM), which did not produce a fluorescent signal. After addition of Casp3 (10 nM) a fluorescent signal was observed with the fastest cleavage rate occurring with CatL. (**b**) Same experiment as (**a**) but the order of protease addition was reversed with first incubation of Casp3 followed by CatL. Casp3 alone did not produce a fluorescent signal compared to sequential addition. (**c**) Negative controls **(D)-Asp 1** and **(D)-Phe 1** do not produce a fluorescent signal after sequential addition of Casp3 followed by CatL, compared to **AND-Gate 1**, which does. (**d**) Same experiment as (**c**) but reversing the order of protease addition. (**e**) Fluorogenic substrate assay with **AND-Gate 1** and respective negative controls. Probes were incubated in tumor lysate derived from excised 4T1 tumors in Balb/C mice.

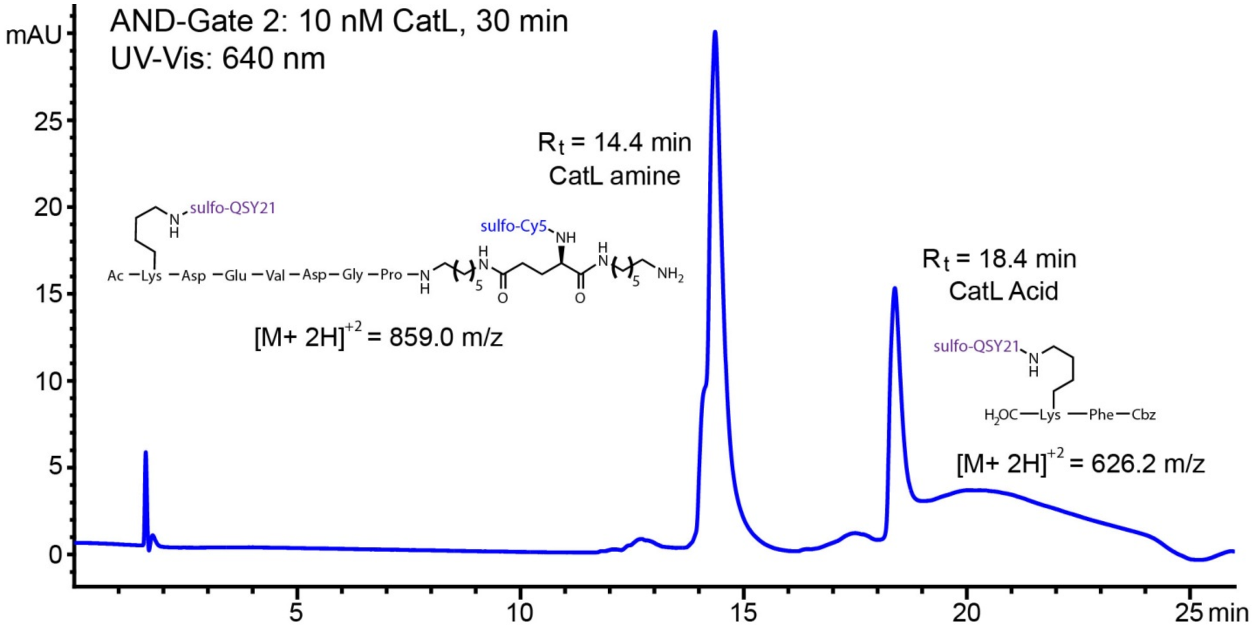
AND-Gate 2: 10 nM CatL, 30 min
UV-Vis: 640 nm
mAU
R_t = 14.4 min
CatL amine
sulfo-QSY21
sulfo-Cy5
Ac –Lys –Asp –Glu –Val –Asp– Gly –Pro
[M+ 2H]^+2 = 859.0 m/z
R_t = 18.4 min
CatL Acid
H2OC—Lys —Phe—Cbz
[M+ 2H]^+2 = 626.2 m/z
min

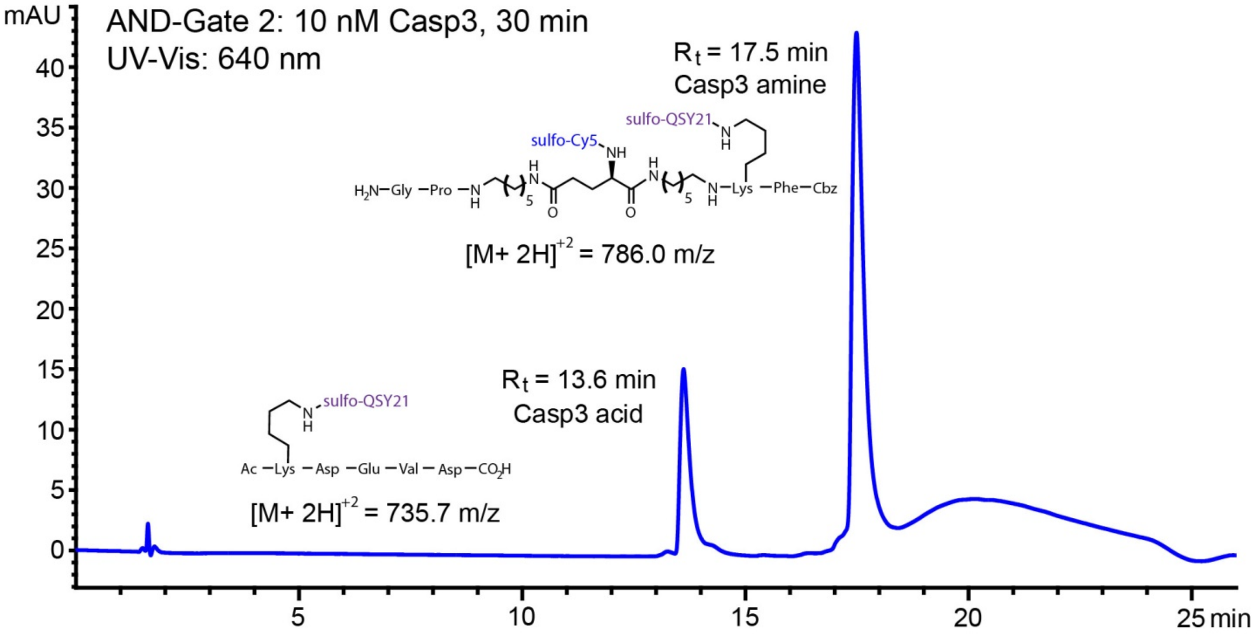
AND-Gate 2: 10 nM Casp3, 30 min
UV-Vis: 640 nm
mAU
R_t = 17.5 min
Casp3 amine
sulfo-QSY21
sulfo-Cy5
H2N–Gly –Pro
Lys –Phe–Cbz
[M+ 2H]^+2 = 786.0 m/z
R_t = 13.6 min
Casp3 acid
Ac –Lys –Asp –Glu –Val – Asp–CO2H
[M+ 2H]^+2 = 735.7 m/z
min

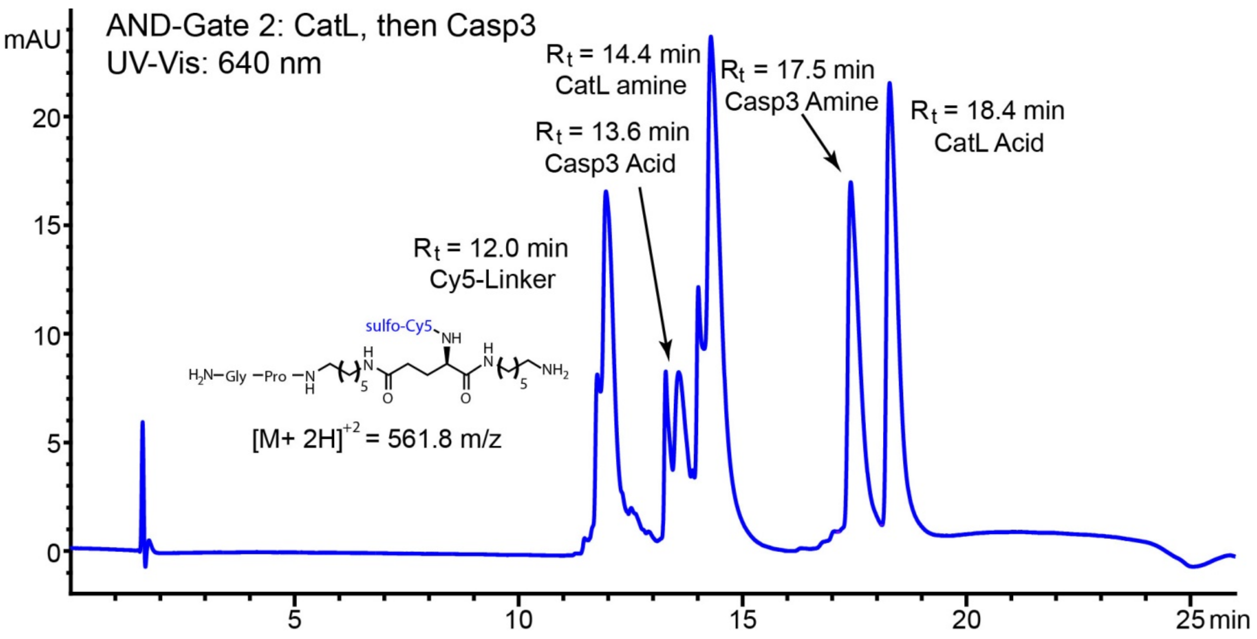


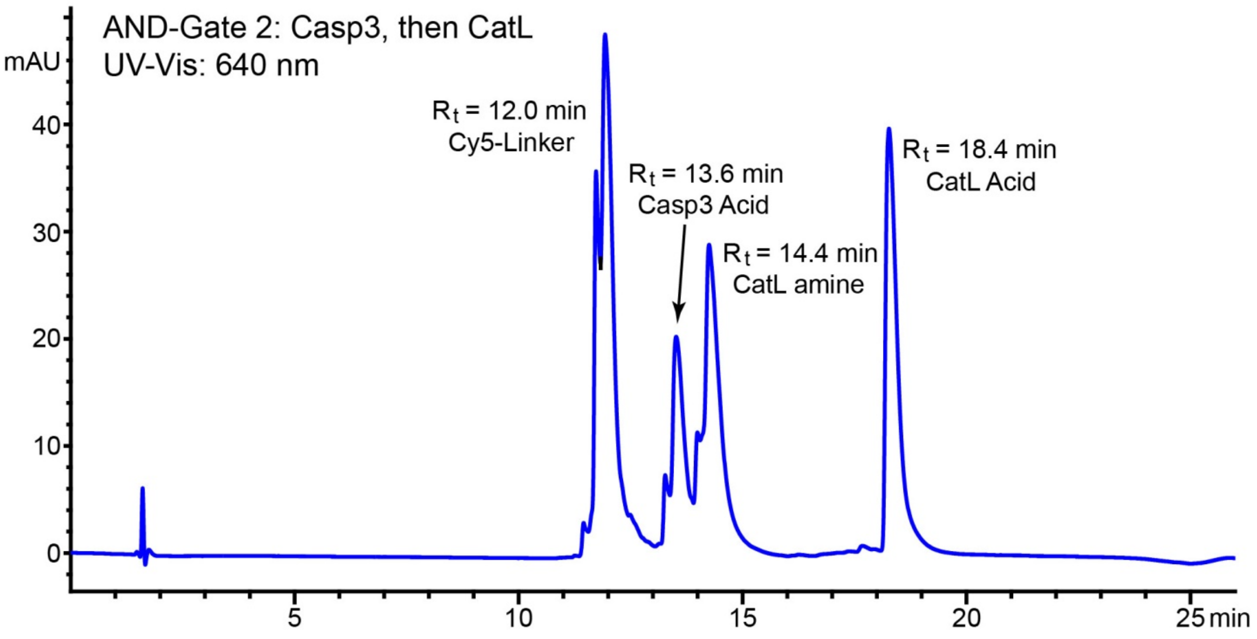


**Supplemental Figure 2.** Detection of **AND-Gate 2** cleavage products using LC-MS. **AND-Gate 2** was incubated with human recombinant CatL or Casp3 alone (10 nM) for 30 min, and with both proteases successively in either order. Structures and the respective charge-to-mass ratios (m/z) of the cleavage products are shown adjacent to the corresponding peaks (wavelength monitored: 640 nm).

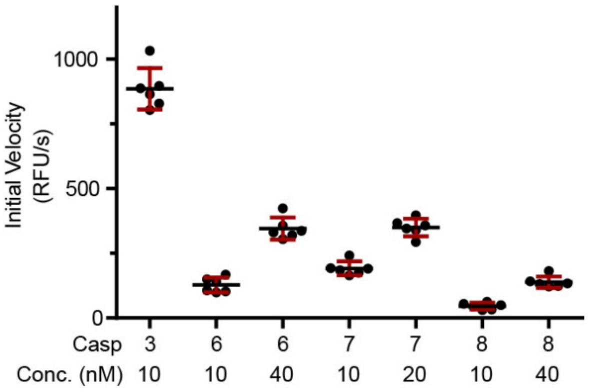


**Supplemental Figure 3.** Evaluation of **AND-Gate 2** with recombinant human caspases using a fluorogenic substrate assay. **AND-Gate 2** was incubated with CatL for 2 h, the assay buffer pH was adjusted to 7.4, and then the respective caspase was added at the indicated concentrations. Initial velocities were calculated from the linear portion of the progress curves for each experiment.

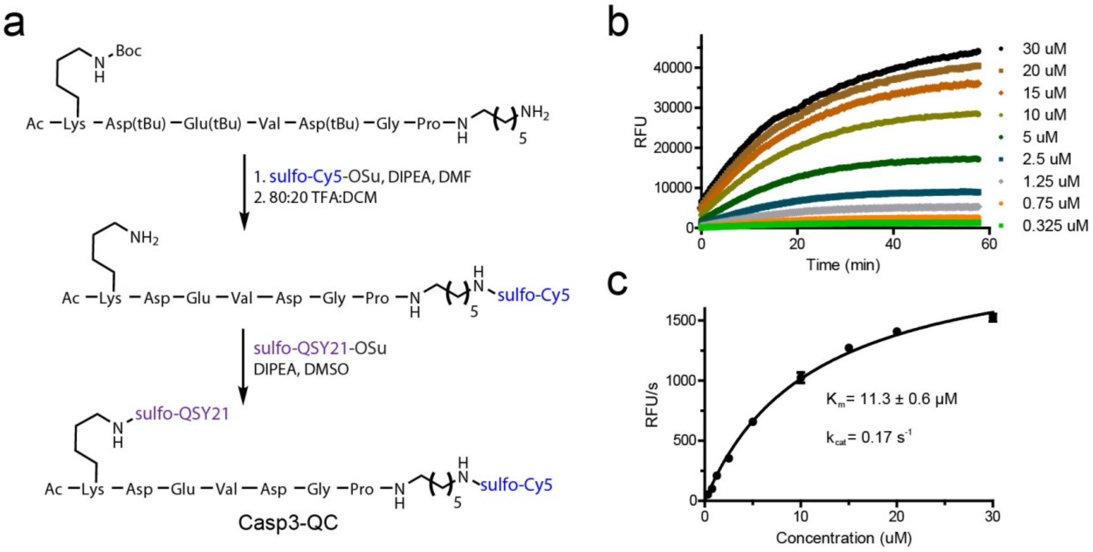


**Supplemental Figure 4.** Synthesis and Characterization of Casp3-QC. (**a**) Synthesis of Casp3-QC probe. For synthetic details see the Methods section below. (**b**) Progress curves of Casp3-QC incubated with Casp3 (10 nM) at various concentrations for calculation of Michaelis-Menten parameters. (**c**) Curve fit of initial velocity versus substrate concentration for Michaelis-Menten parameters.

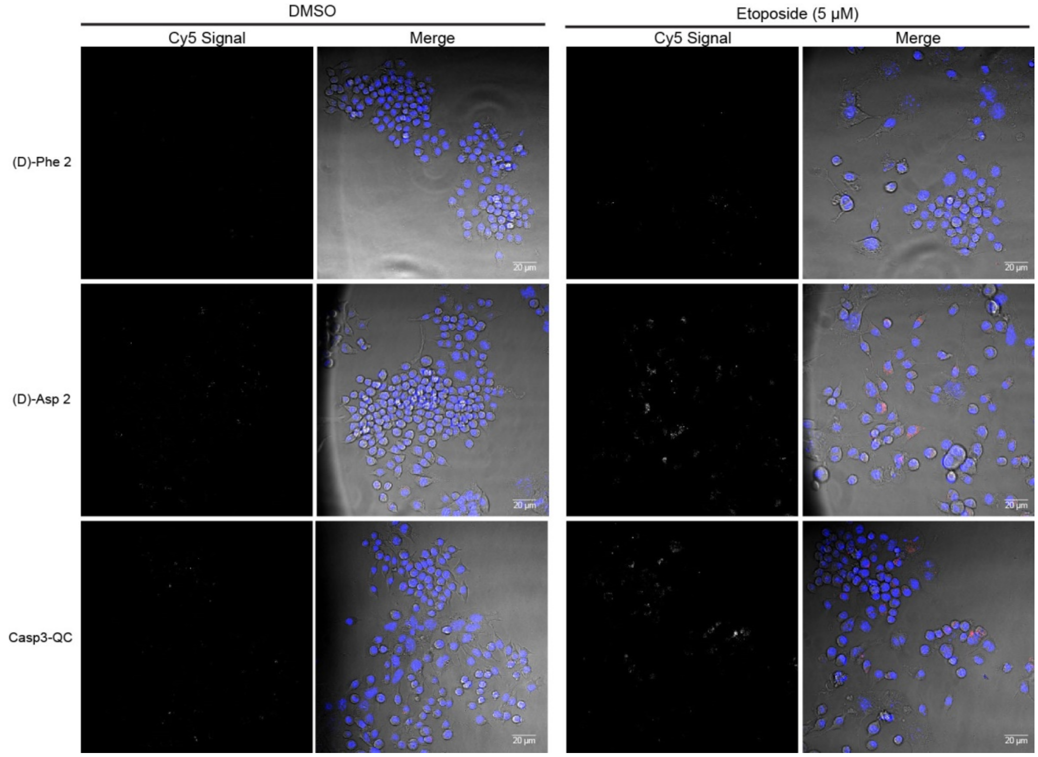


**Supplemental Figure 5.** Representative microscopy images of negative controls **(D)Phe 2**, **(D)Asp 2**, and **Casp3-QC** for the 4T1:RAW macrophage co-culture experiment described in **Figure 3e**. Cy5 signal is shown in grayscale in the left panels. For the merge image: Blue is nuclear staining and red is Cy5 probe signal, brightfield is in grayscale. All images are normalized.

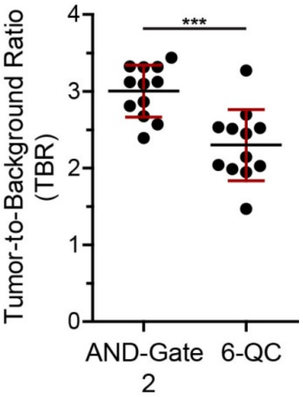


**Supplemental Figure 6.** Tumor-to-background ratio (TBR) of **AND-Gate 2** and **6-QC** in 4T1 tumors compared to adjacent healthy fat tissue. The area used to measure fluorescent signal within the tumor and adjacent tissue was normalized across all samples.

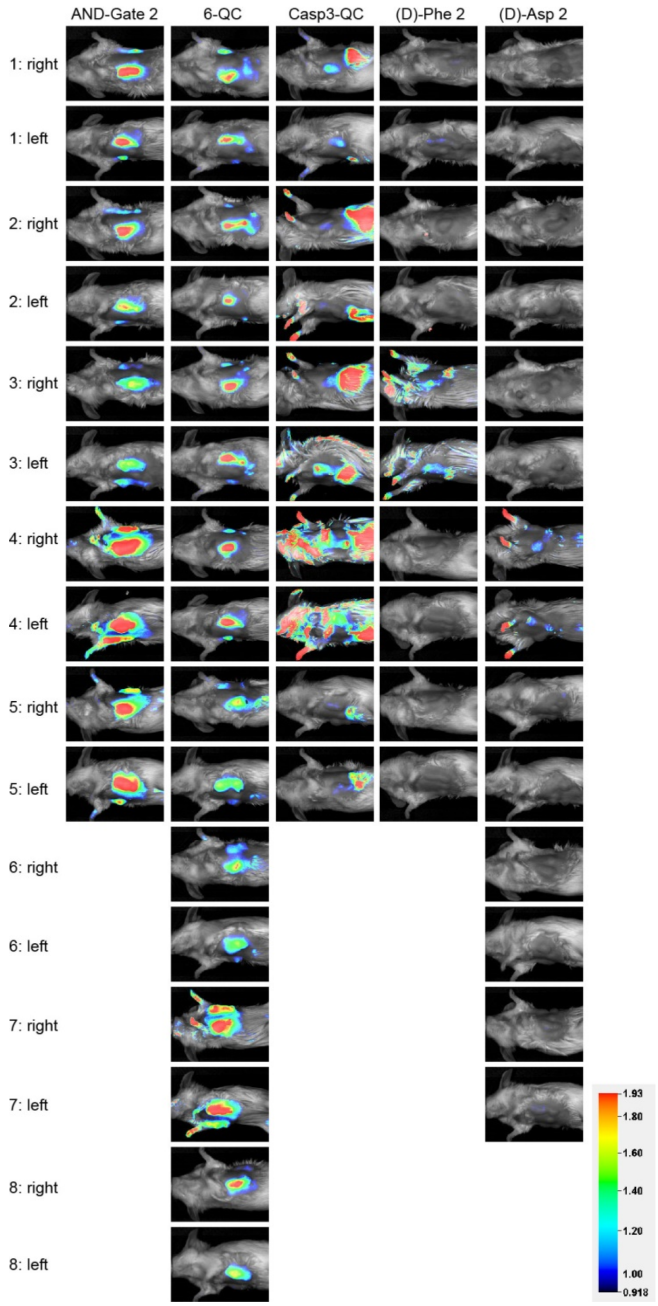


**Supplemental Figure 7.** Rainbow plots of fluorescent signal in 4T1 breast tumor bearing Balb/C mice. Images are of the left and right tumors for each mouse respectively. The fluorescent signal in all images are normalized. Each probe was injected 2 h prior to imaging via I.V. tail vein (20 nmol). For experimental details see the Biology Methods section.

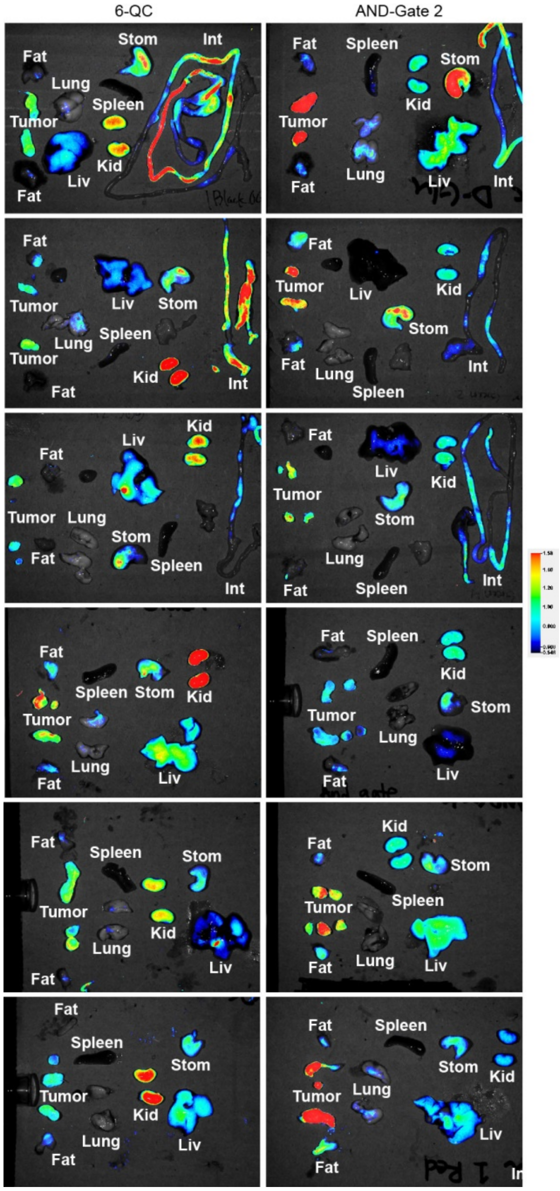


**Supplemental Figure 8.** Rainbow plots of excised tumor and tissues from Balb/C mice bearing 4T1 tumors 2 h post I.V. tail vein injection of **6-QC** or **AND-Gate 2** (20 nmol).

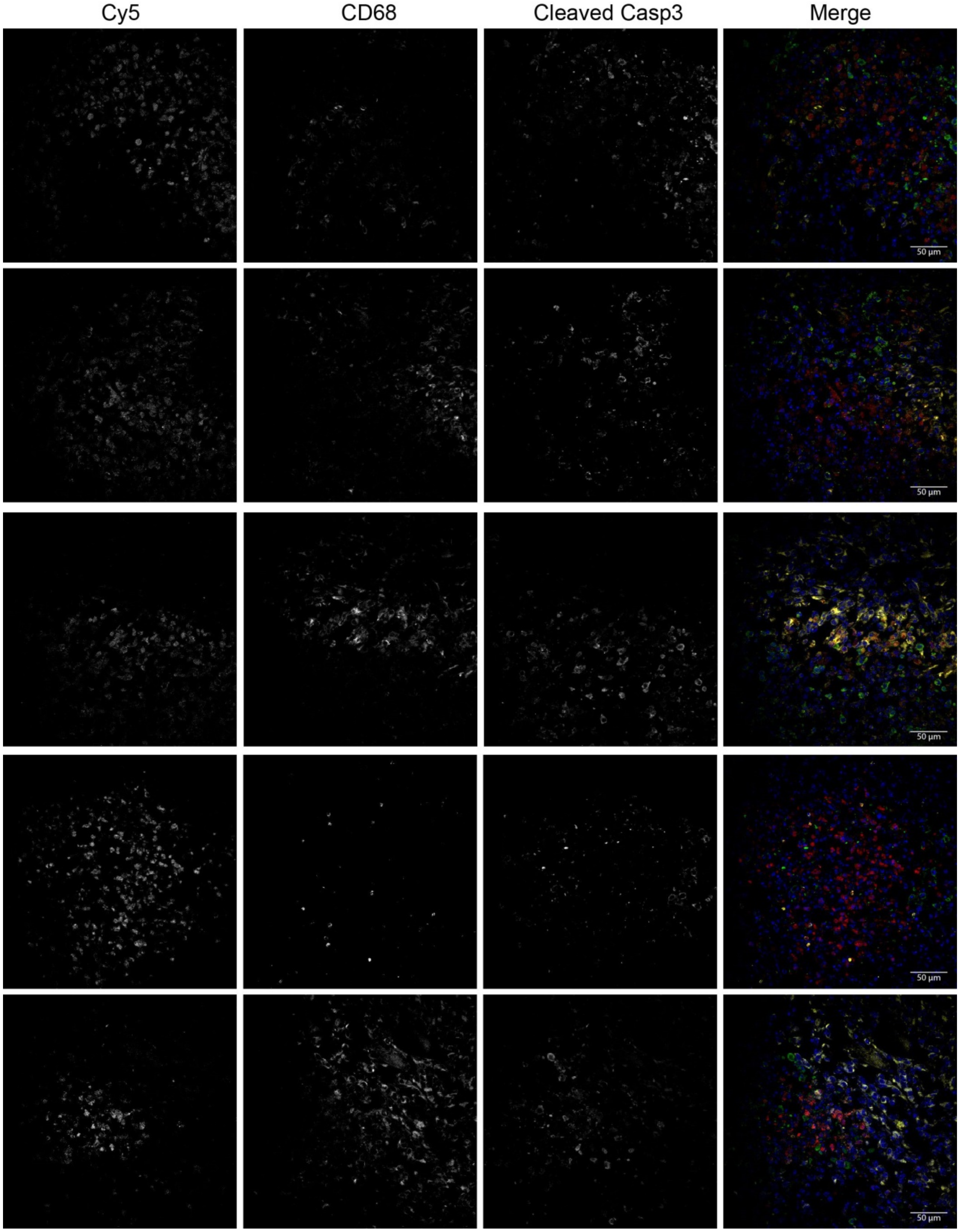


**Supplemental Figure 9.** Additional example images of sectioned 4T1 breast tumor tissue from mice injected with **AND-Gate 2** and stained for CD68 (macrophage marker) and cleaved Casp3. The Cy5 channel corresponds to the fluorescent signal from probe. All single channel images are in gray scale. For merged images: Cy5 = Red, CD68 = Yellow, Cleaved Casp3 = Green, DAPI = Blue.

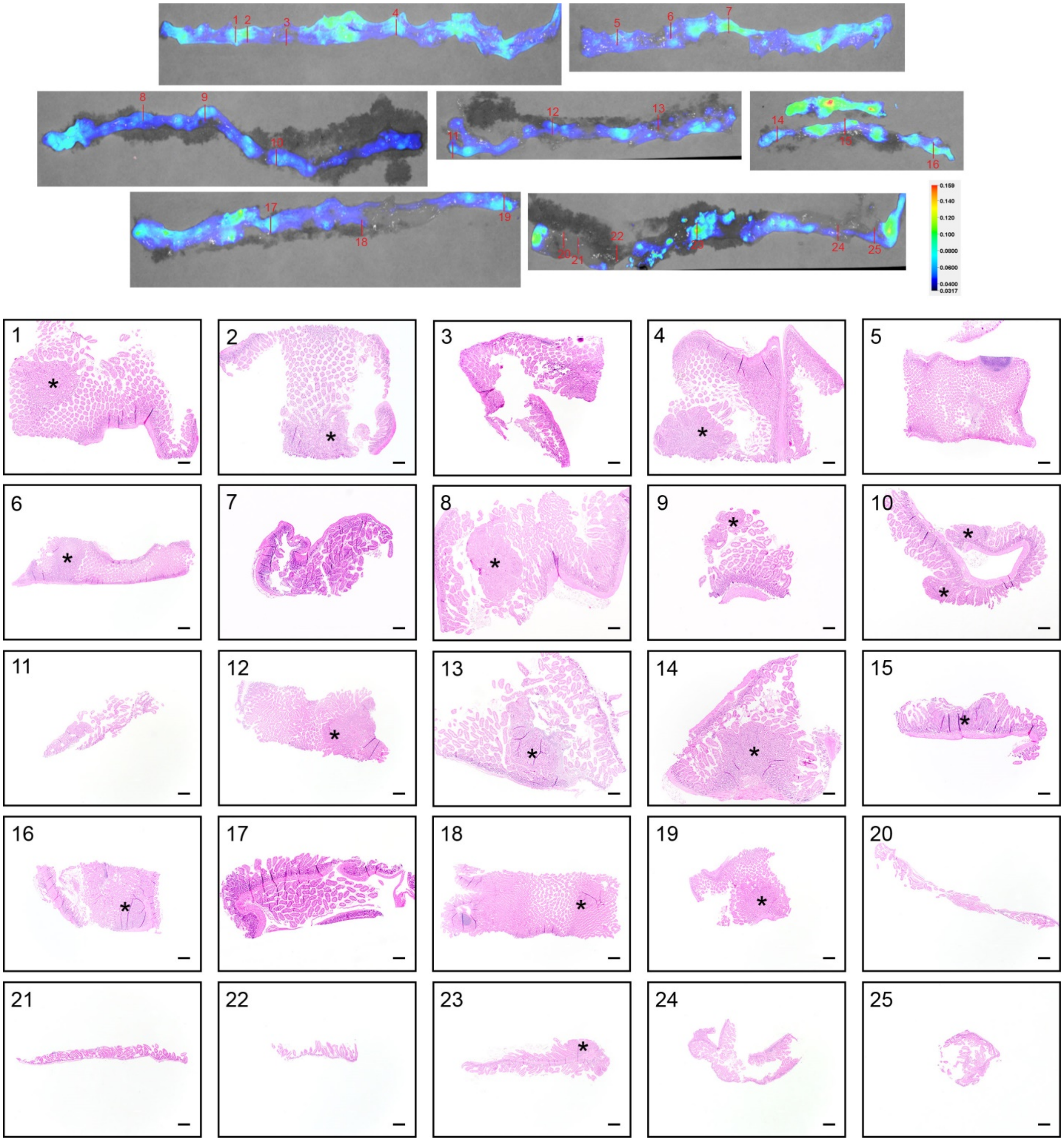

**Supplemental Figure 10.** Rainbow plots of splayed intestines excised from mice injected with **6-QC** (R.O., 20 nmol) with indicated positions taken for sectioning and evaluated for the presence of tumor by H&E staining. Fluorescent signal is normalized between all images. Scale bars are 20 μm.

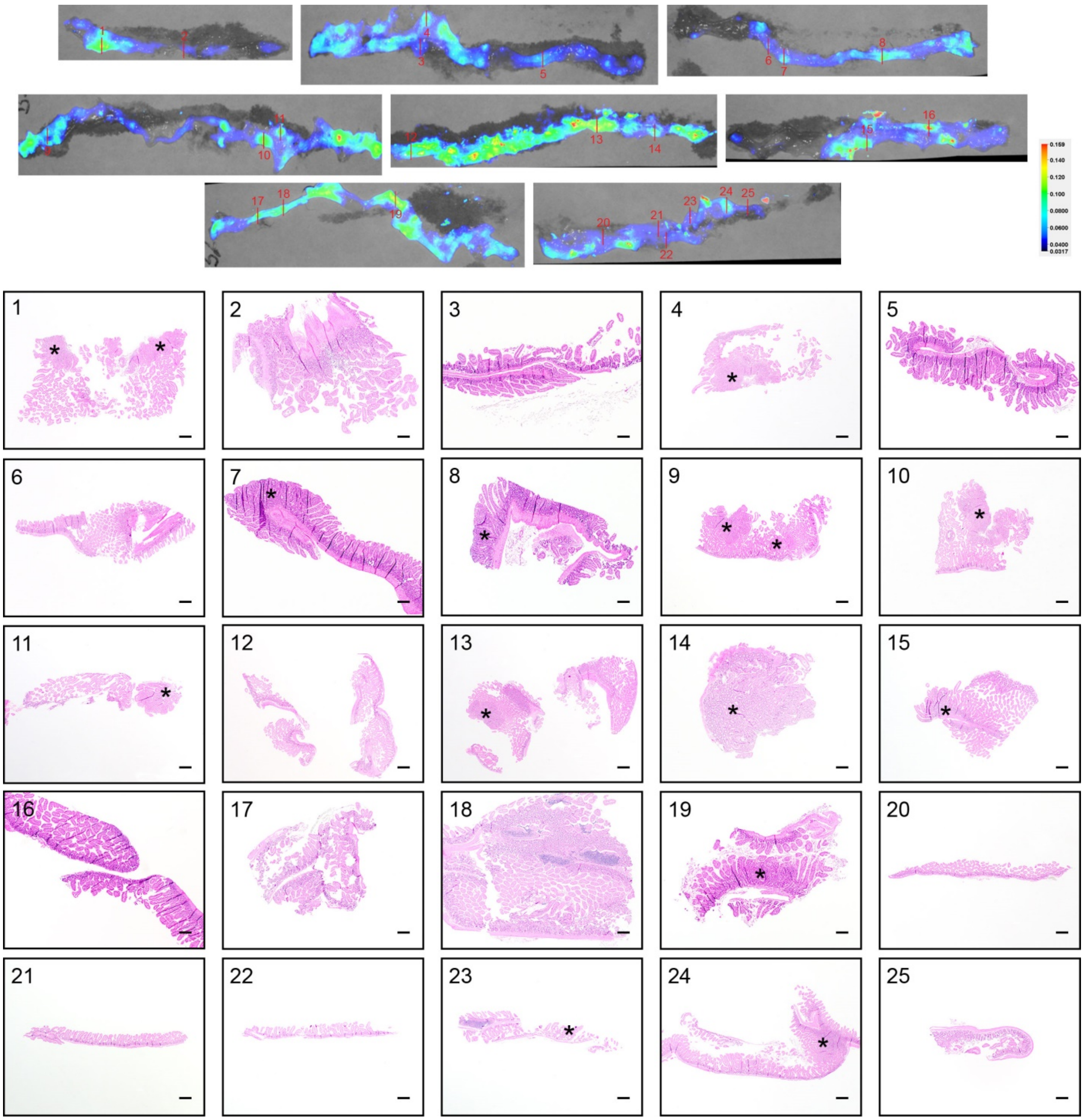

**Supplemental Figure 11.** Rainbow plots of splayed intestines excised from mice injected with **AND-Gate 2** (R.O., 20 nmol) with indicated positions taken for sectioning and evaluated for the presence of tumor by H&E staining. Fluorescent signal is normalized between all images. Scale bars are 20 μm.

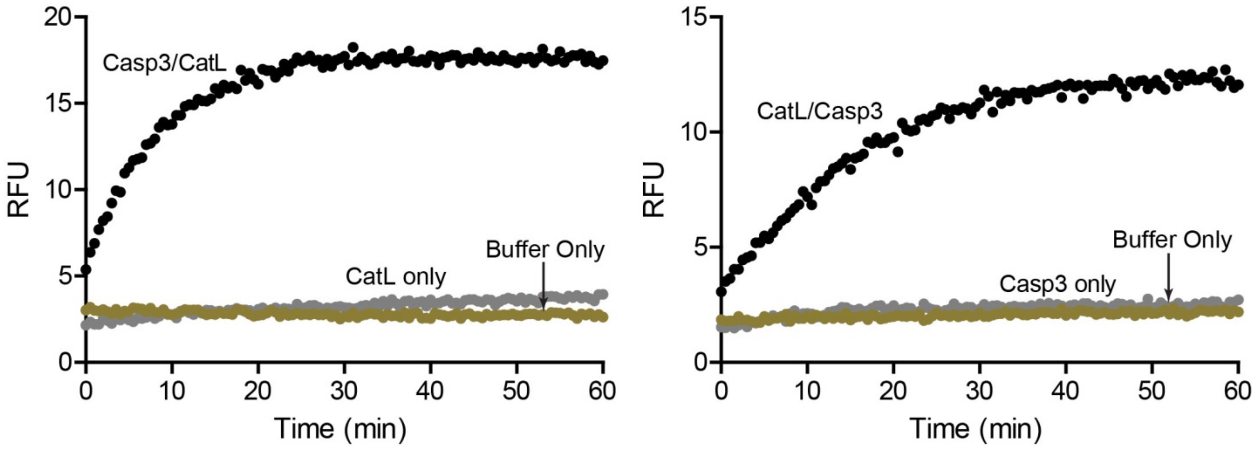


**Supplemental Figure 12**. Fluorogenic substrate assays with **AND-Gate-FNIR**. Probe was incubated with either protease first followed by addition of the second protease. The probe is fluorescently activated only after incubation with both proteases regardless of order of addition whereas addition of a single protease or buffer does not produce a signal.

### III. Chemistry Methods

**Materials and Synthetic Methods.** All reactions were performed exposed to atmospheric air and with solvents not previously dried over molecular sieves or other drying agents. Reactions containing light sensitive materials were protected from light. The ACS reagent grade *N*,*N*'-dimethylformamide (DMF), tetrahydrofuran (THF) containing 250 ppm of butylated hydroxy toluene (BHT), molecular biology grade dimethyl sulfoxide (DMSO), and all other commercially available chemicals were used without further purification. Reaction temperatures above 23 °C refer to incubator temperatures, controlled by a temperature modulator. Reaction progress and purity analysis was monitored using an analytical LC-MS. The LC-MS systems used was either a Thermo Fisher Finnigan Surveyor Plus equipped with an Agilent Zorbax 300SB-$C_{18}$ column (3.5 µm, 3.0 x 150 mm) coupled to a Finnigan LTQ mass spectrometer or an Agilent 1100 Series HPLC equipped with a Luna 4251-E0 $C_{18}$ column (3 µm, 4.6 x 150 mm) coupled to a PE SCIEX API 150EX mass spectrometer (wavelengths monitored = 215 & 254 nm). Purification of intermediates and final compounds was carried out using either a semi-preparative Luna $C_{18}$ column (5 µm, 10 x 250 mm) attached to an Agilent 1260 Infinity HPLC system or a CombiFlash Companion/TS (Teledyne Isco) with a 4 or 12 g reverse phase $C_{18}$ RediSep Rf Gold column (wavelengths monitored = 215 & 254 nm). Information regarding gradient programs for purifications can be found in the Chemistry Protocols section below. Intermediates were identified by their expected m/z using LC-MS or direct injection MS.

### Chemistry Protocols.

Solid Phase Peptide Synthesis. The Cat (**1**), Casp3 (**3**) and negative control substrates were synthesized on 2-Chlorotrityl resin using standard Fmoc chemistry as previously described[2]. Peptides were cleaved from resin using 1,1,1,2,2,2-hexafluoroisopropanol to maintain the protecting groups on the amino acid side chains[3]. All peptides were reverse phase HPLC purified and lyophilized prior to use.

General Procedure A: Amide bond Coupling. The coupling reagent O-(1H-6-Chlorobenzotriazole-1-yl)-1,1,3,3-tetramethyluronium hexafluorophosphate (HCTU) was dissolved with the carboxylic acid starting material and 2,4,6-collidene in DMF. The solution of activated acid was added to the amine and agitated at RT.

General Procedure B: Allyl Deprotection. $Pd(OAc)_2$ (0.5 equiv) was mixed with triphenylphosphine ($PPh_3$, 1 equiv) in THF. The solution of activated $Pd^0$ was added to the allyl starting material (1 equiv). Then, phenylsilane ($SiPhH_3$, 5 equiv) was added to the solution and stirred at RT for 16 h. After the reaction, the solution was concentrated *in vacuo* and dissolved in 1:1 MeCN:$H_2O$ (0.1% TFA) for purification via reverse phase chromatography.

General Procedure C: Boc and tBu peptide sidechain deprotection. After the respective amide bond coupling and HPLC purification, the purified product was collected and concentrated *in vacuo*. The product, was then dissolved 7:2:0.5:0.5 TFA:DCM:$H_2O$:TIS and stirred at RT for 2 h. The reaction was then concentrated *in vacuo*. The residue was dissolved in 1:1 MeCN:$H_2O$ (0.1% TFA) and lyophilized.

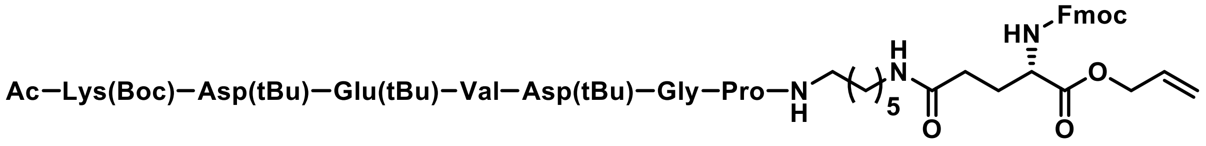


**Intermediate 1**. General Procedure A–**S1** (17 mg, 0.015 mmol), Fmoc-Glu(OH)-OAll (12 mg, 0.029 mmol), HCTU (12 mg, 0.029 mmol), 2,4,6-collidene (7.5 µL, 0.058 mmol), DMF (500 µL). The reaction was stirred for 16 h and then concentrated *in vacuo*. The reaction was dissolved in 7:3 MeCN:$H_2O$ (0.1% TFA) and purified using semi-prep reverse HPLC with a gradient program of 10% MeCN:$H_2O$ (0.1% TFA) for 0-2 min, 10-95% for 2-22 min, 95% for 22-26 min ($R_t$ = 18.5 min). The purified fractions were collected and lyophilized to obtain a white powder (11 mg, 45% yield).

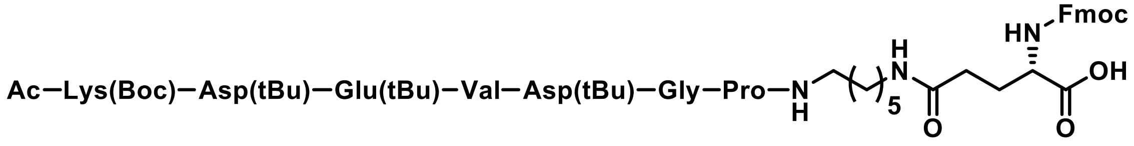


**S1**. General Procedure B–**Intermediate 1** (11 mg, $7.1x10^{-3}$ mmol), $Pd(OAc)_2$ (1 mg, $3.5x10^{-3}$ mmol), $PPh_3$ (2 mg, $7.1x10^{-3}$ mmol), $PhSiH_3$ (2.6 µL, 0.021 mmol), THF (1 mL). The reaction was purified using semi-prep reverse phase HPLC with a gradient of 2% MeCN:$H_2O$ (0.1% TFA) for 0-2 min, 2-75% for 2-23 min, 75-95% for 23-26 min ($R_t$ = 22.1 min). The purified fractions were collected and lyophilized to obtain a white powder (7 mg, 63% yield).

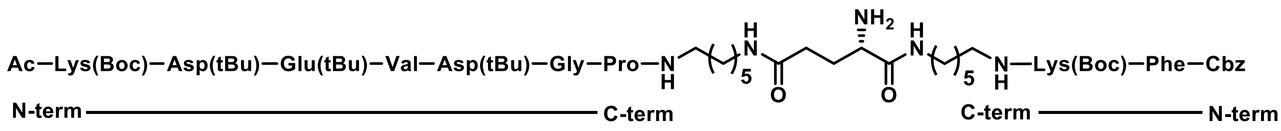


**Intermediate 2**. General Procedure A–**S1** (7 mg, 4.5x10$^{-3}$ mmol), Cat substrate **1** (3 mg, 4.5x10$^{-3}$ mmol), HCTU (6 mg, 1.4x10$^{-2}$ mmol), 2,4,6-collidene (2.9 μL, 2.3x10$^{-3}$ mmol), DMF (500 μL). The reaction was agitated for 16 h followed by addition of piperidine (120 μL) and then agitated for an additional 1 h. Then, the reaction was concentrated *in vacuo*. The reaction was dissolved in 1:1 MeCN:$H_2O$ (0.1% TFA) and purified using semi-prep reverse HPLC with a gradient of 2% MeCN:$H_2O$ (0.1% TFA) for 0-2 min, 2-75% for 2-23 min, 75-95% for 23-26 min ($R_t$ = 19 min). The purified fractions were collected and lyophilized to obtain a white powder (5 mg, 55% yield).

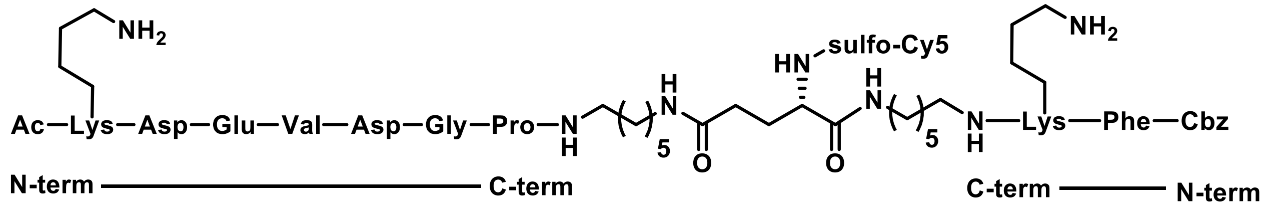


**4**. **Intermediate 2** (5 mg, 2.5x10$^{-3}$ mmol) and sulfo-Cy5-OSu (4 mg, 5.0x10$^{-3}$ mmol) were dissolved in DMF (500 μL). Then, DIPEA (2.2 μL, 1.3x10$^{-2}$ mmol) was added and the reaction was agitated for 24 h at RT. The reaction was quenched with the addition of 1:1 MeCN:$H_2O$ (0.1% TFA) and purified using semi-prep reverse HPLC with a gradient of 2% MeCN:$H_2O$ (0.1% TFA) for 0-2 min, 2-75% for 2-23 min, 75-95% for 23-26 min ($R_t$ = 22.8 min). The purified fractions were collected and lyophilized to obtain a white powder (5 mg, 55% yield). The purified fractions were collected and General Procedure C was followed to obtain a blue powder (no yield given).

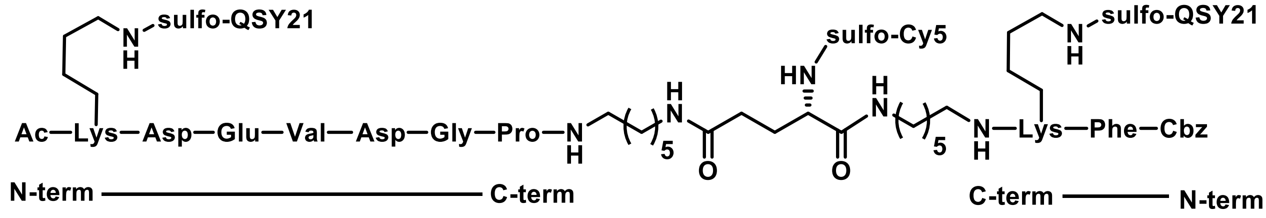


**AND-Gate 1**. Intermediate **S4** (5.4 mg, 2.5x10$^{-3}$ mmol) and sulfo-QSY21-OSu (7.1 mg, 1.0x10$^{-2}$ mmol) were dissolved in DMSO (600 μL). Then, DIPEA was added (2.2 μL, 1.3x10$^{-2}$ mmol) and the reaction was agitated for 24 h at 37 °C. The reaction was quenched by the addition of 1:1 MeCN:H2O (0.1% TFA) and purified with reverse phase semi-prep HPLC with a gradient program of 2% MeCN:H2O (0.1% TFA) for 0-2 min, 2-40% for 2-32 min, 40-95% for 32-36 min ($R_t$ = 31.7 min). Purified fractions were collected and lyophilized to obtain a blue powder (2.39 mg, 25% yield over last 2 steps). ESI-HRMS (m/z) calculated for $C_{187}H_{218}N_{24}O_{47}S_8^{2+}$: 1904.6614; found 1904.6579.

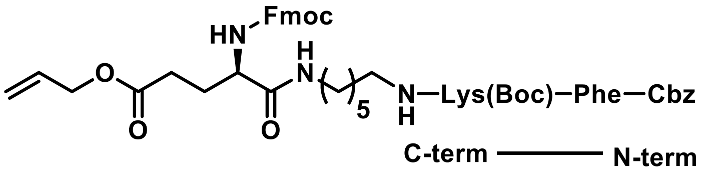


**Intermediate 3**. General Procedure A–**S2** (23 mg, 0.036 mmol), Fmoc-Glu(OAll)-OH (44 mg, 0.108 mmol), HCTU (47 mg, 0.108 mmol), 2,4,6-collidene (23 µL, 0.18 mmol), DMF (1 mL). The reaction was stirred for 16 h and then concentrated *in vacuo*. The products were dissolved in 7:3 MeCN:$H_2O$ (0.1% TFA) and purified using a reverse phase Combiflash with a gradient program of 10% MeCN:$H_2O$ (0.1% TFA) for 0-2 min, 10-80% for 2-19 min ($R_t$ = 15.5 min). The purified fractions were collected and lyophilized to obtain a white powder (29 mg, 80% yield).

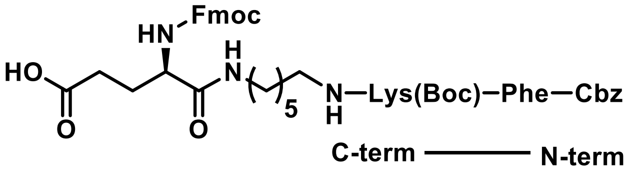


**2**. General Procedure B–**Intermediate 3** (29 mg, 0.029 mmol), $Pd(OAc)_2$ (3 mg, 0.014 mmol), $PPh_3$ (11 mg, 0.043 mmol), $PhSiH_3$ (10 µL, 0.086 mmol), THF (1 mL). The reaction was purified using a reverse phase Combiflash with a gradient of 10% MeCN:$H_2O$ (0.1% TFA) for 0-2 min, 10-80% for 2-19 min, 80-95% for 19-25 min ($R_t$ = 16 min). The purified fractions were collected and lyophilized to obtain a white powder (27 mg, 95% yield).

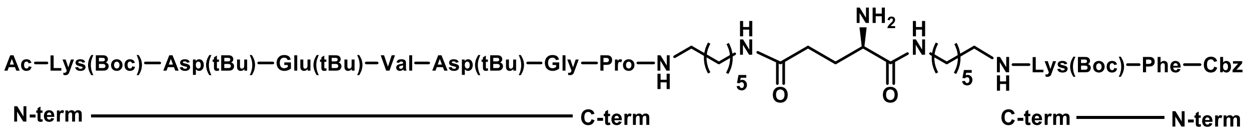


**Intermediate 4**. General Procedure A–**2** (11 mg, 0.011 mmol), Casp3 substrate **3** (19 mg, 0.016 mmol), HCTU (5 mg, 0.011 mmol), 2,4,6-collidene (4.2 µL, 0.032 mmol), DMF (500 µL). The reaction was agitated for 16 h followed by addition of piperidine (120 µL) and then agitated for an additional 1 h. Then, the reaction was concentrated *in vacuo*. The reaction was dissolved in 1:1 MeCN:$H_2O$ (0.1% TFA) and purified using a reverse phase Combiflash (4 g column) with a gradient program of 10% MeCN:$H_2O$ (0.1% TFA) for 0-2 min, 10-70% for 2-20 min ($R_t$ = 17.2 min). The purified fractions were collected and lyophilized to obtain a white powder (8 mg, 40% yield).

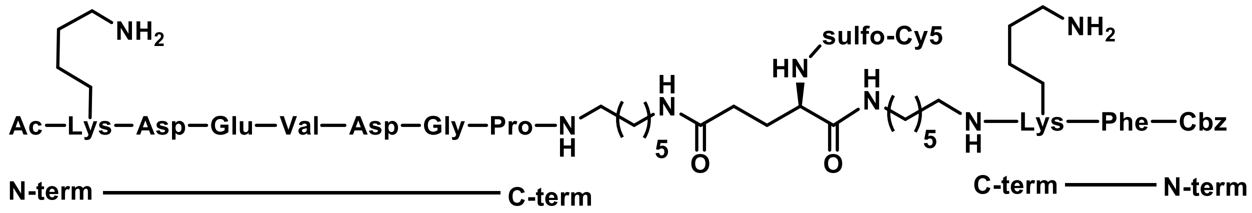


4. **Intermediate 2** (8 mg, 4.3x10$^{-3}$ mmol) and sulfo-Cy5-OSu (5 mg, 6.5x10$^{-3}$ mmol) were dissolved in DMF (600 µL). Then, DIPEA (3.8 µL, 2.2x10$^{-2}$ mmol) was added and the reaction was agitated for 24 h at RT. The reaction was quenched with the addition of 1:1 MeCN:$H_2O$ (0.1% TFA) and purified using semi-prep reverse HPLC with a gradient program of 10% MeCN:$H_2O$ (0.1% TFA) for 0-2 min, 10-95% for 2-23 min, 95% for 23-26 min ($R_t$ = 18.6 min). The purified fractions were collected and General Procedure C was followed to obtain a blue powder (no yield given).

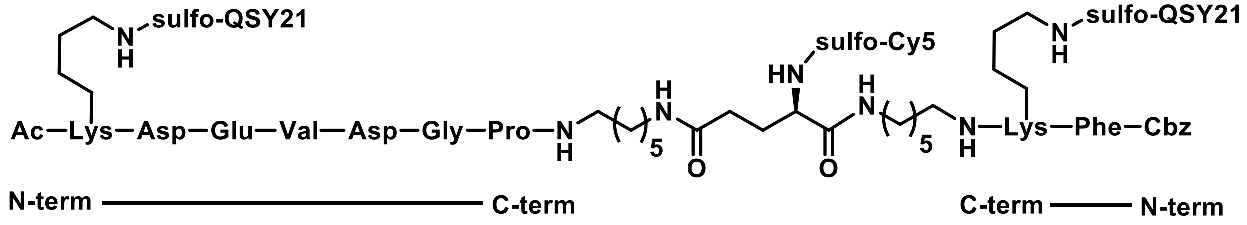


**AND-Gate 2**. Compound **4** (5.9 mg, 2.7x10$^{-3}$ mmol) was combined with sulfo-QSY21-OSu (6.4 mg, 6.09x10$^{-3}$ mmol) and then dissolved in DMSO (600 µL). Then, DIPEA was added (2.4 µL, 1.4x10$^{-2}$ mmol) and the reaction was agitated for 24 h at 37 °C. The reaction was quenched by the addition of 1:1 MeCN:H2O (0.1% TFA) and purified with reverse phase semi-prep HPLC with a gradient program of 5% MeCN:H2O (0.1% TFA) for 0-2 min, 5-40% for 2-32 min, 40-95% for 32-36 min ($R_t$ = 31.8 min). Purified fractions were collected and lyophilized to obtain a blue powder (3.5 mg, 34% yield). ESI-HRMS (m/z) calculated for $C_{187}H_{219}N_{24}O_{47}S_8^{3+}$: 1270.1100; found 1270.1087.

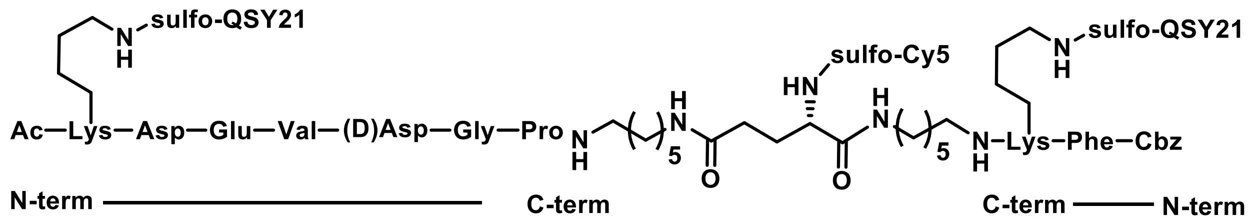


**(D)-Asp 1**. The negative control molecule **(D)-Asp 1** was synthesized in the same manner as **AND-Gate 1** with no significant changes to yields, retention times, or purification protocols. Overall synthesis yield, 7%. ESI-HRMS (m/z) calculated for $C_{187}H_{219}N_{24}O_{47}S_8^{3+}$: 1270.1100; found 1270.1085.

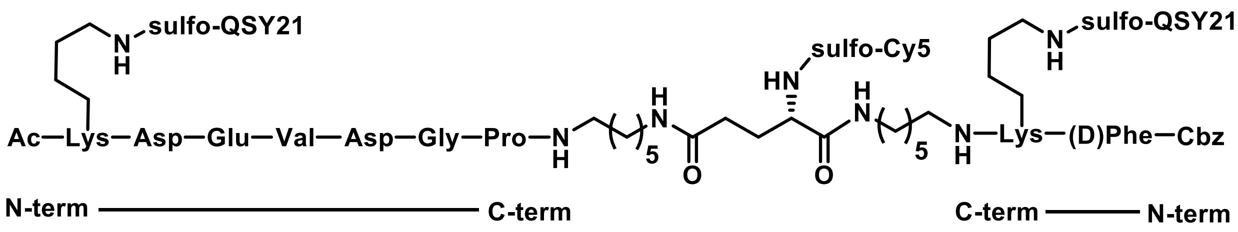


**(D)-Phe 1**. The negative control molecule **(D)-Phe 1** was synthesized in the same manner as **AND-Gate 1** with no significant changes to yields, retention times, or purification protocols. Overall synthesis yield, 12%. ESI-HRMS (m/z) calculated for $C_{187}H_{219}N_{24}O_{47}S_8^{3+}$: 1270.1100; found 1270.1089.

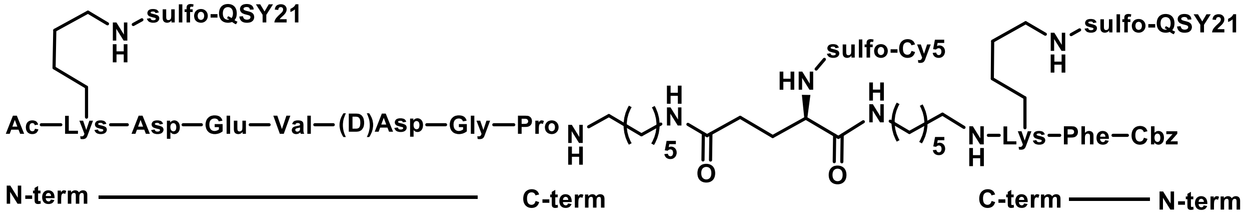


**(D)-Asp 2**. The negative control molecule **(D)-Asp 2** was synthesized in the same manner as **AND-Gate 2** with no significant changes to yields, retention times, or purification protocols. Overall synthesis yield, 3%. ESI-HRMS (m/z) calculated for $C_{187}H_{219}N_{24}O_{47}S_8^{3+}$: 1270.1100; found 1270.1087. ESI-HRMS (m/z) calculated for $C_{187}H_{219}N_{24}O_{47}S_8^{3+}$: 1270.1100; found 1270.1087.

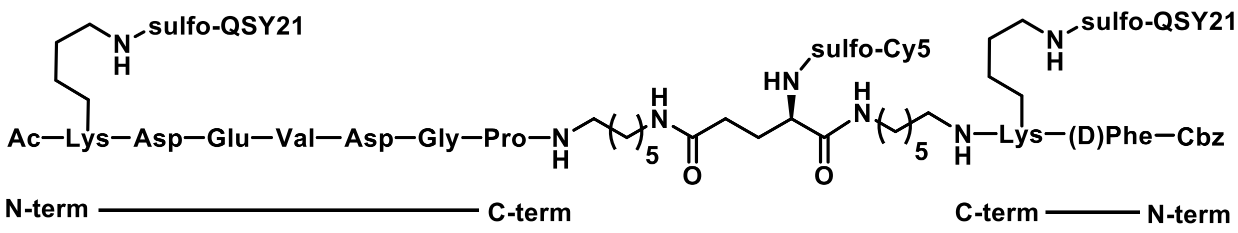


**(D)-Phe 2**. The negative control molecule **(D)-Phe 2** was synthesized in the same manner as **AND-Gate 2** with no significant changes to yields, retention times, or purification protocols. Overall synthesis yield, 4%. ESI-HRMS (m/z) calculated for $C_{187}H_{219}N_{24}O_{47}S_8^{3+}$: 1270.1100; found 1270.1105.

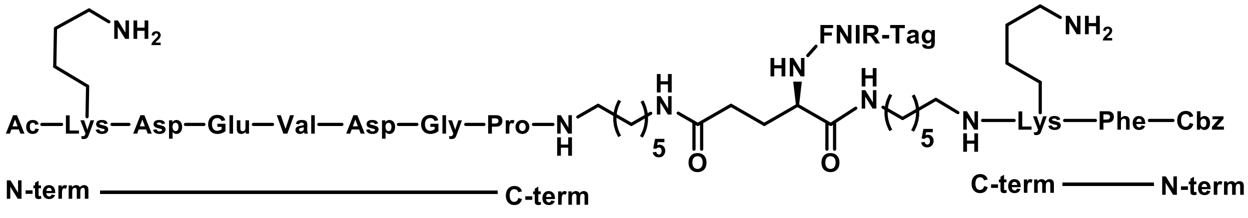

**S2**. **Intermediate 2** (2.9 mg, 1.5x10$^{-3}$ mmol) and FNIR-Tag-OSu (3.5 mg, 3.0x10$^{-3}$ mmol) were dissolved in DMF (50 µL). Then, DIPEA (3.0 µL, 1.5x10$^{-2}$ mmol) was added and the reaction was agitated for 24 h at RT. The reaction was quenched with the addition of 1:1 MeCN:$H_2O$ (0.1% TFA) and purified using semi-prep reverse HPLC with a gradient program of 20% MeCN:$H_2O$ (0.1% TFA) for 0-2 min, 20-70% for 2-20 min, 70% for 20-22 min, 70-95% for 22-24 min, 95% for 24-30 min ($R_t$ =21.3 min). The purified fractions were collected and General Procedure C was followed to obtain a dark green powder (2.6 mg, 65% yield).

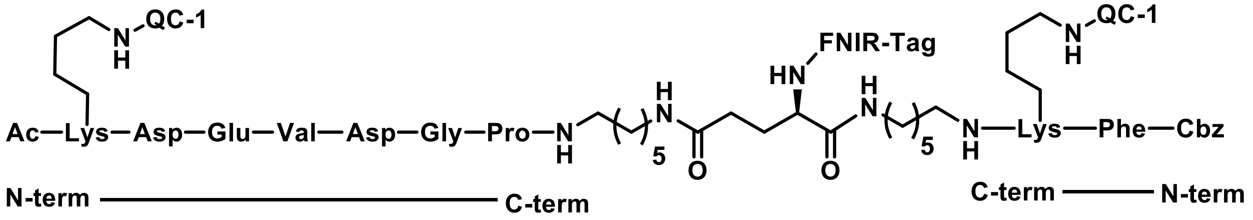


**AND-Gate-FNIR**. **S2** (2.6 mg, 9.8x10$^{-4}$ mmol) was combined with QC1-OSu (5.8 mg, 4.9x10$^{-3}$ mmol) and then dissolved in DMSO (60 µL). Then, DIPEA was added (1.7 µL, 9.8x10$^{-3}$ mmol) and the reaction was agitated for 24 h at 37 °C. The reaction was quenched by the addition of 1:1 MeCN:H2O (0.1% TFA) and purified with reverse phase semi-prep HPLC with a gradient program of 5% MeCN:H2O (0.1% TFA) for 0-2 min, 5-40% for 2-20 min, 40% for 20-23 min ($R_t$ = 21.3 min). Purified fractions were collected and lyophilized to obtain a blue powder (1.9 mg, 41% yield). ESI-HRMS (m/z) calculated for $C_{224}H_{313}Cl_2N_{25}O_{60}S_{10}^{4+}$: 1176.221; found 1175.7230.

### IV. HPLC Purity Analysis

#### AND-Gate 1

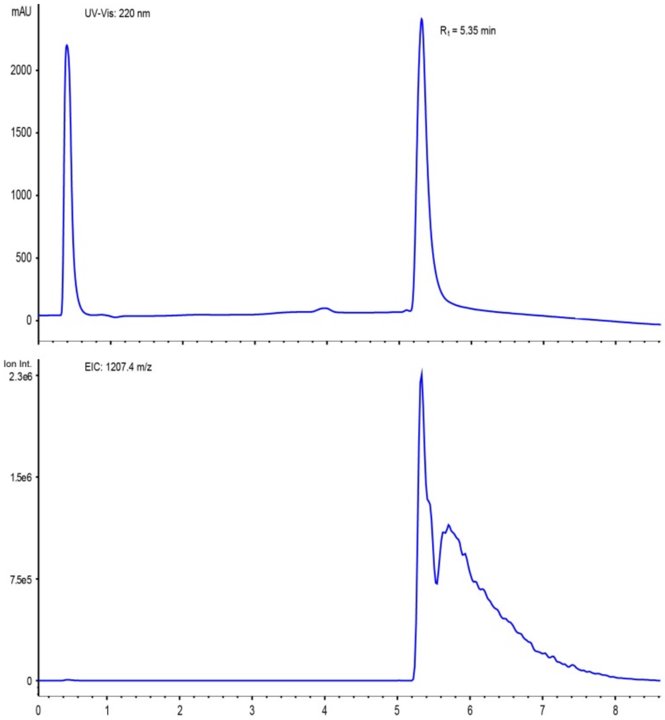


#### (D)-Asp 1

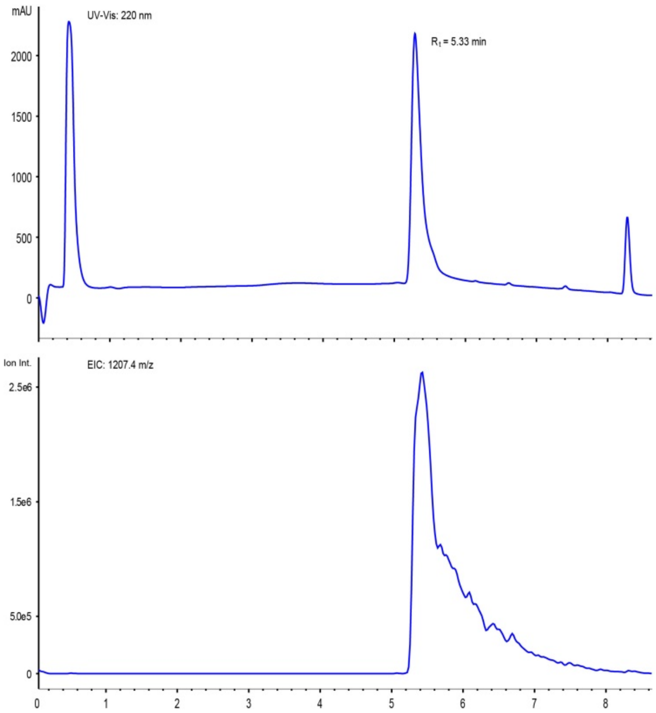

**(D)-Phe 1**

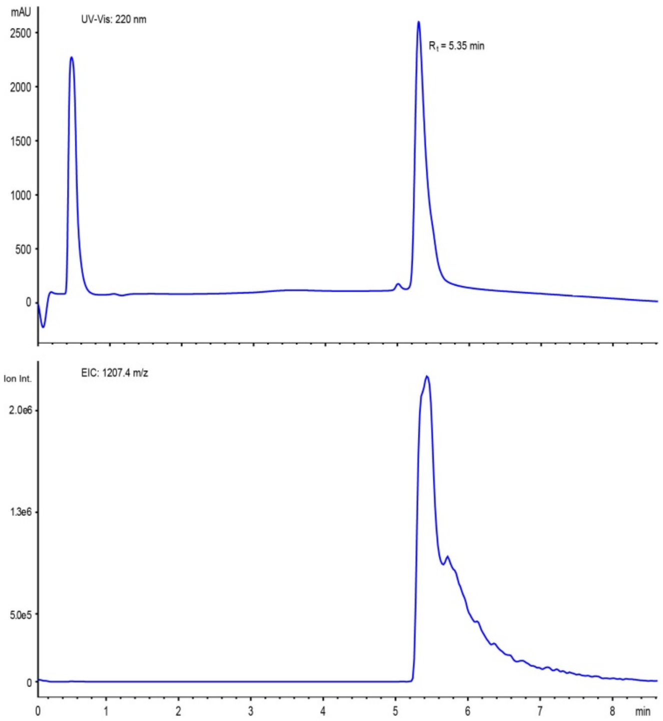


**AND-Gate 2**

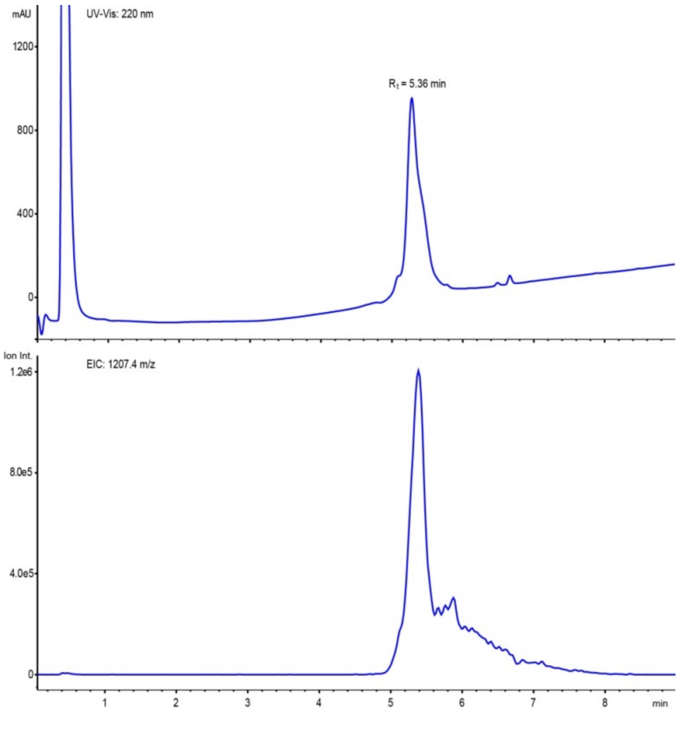

**(D)-Asp 2**

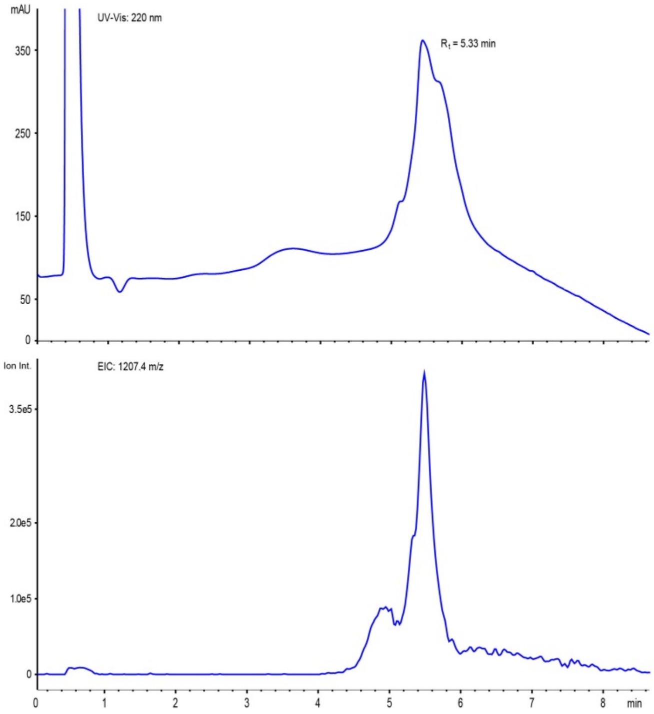
mAU
UV-Vis: 220 nm
$R_t$ = 5.33 min
350
250
150
50
0
Ion Int.
EIC: 1207.4 m/z
3.5e5
2.0e5
1.0e5
0
0
1
2
3
4
5
6
7
8
min

**(D)-Phe 2**

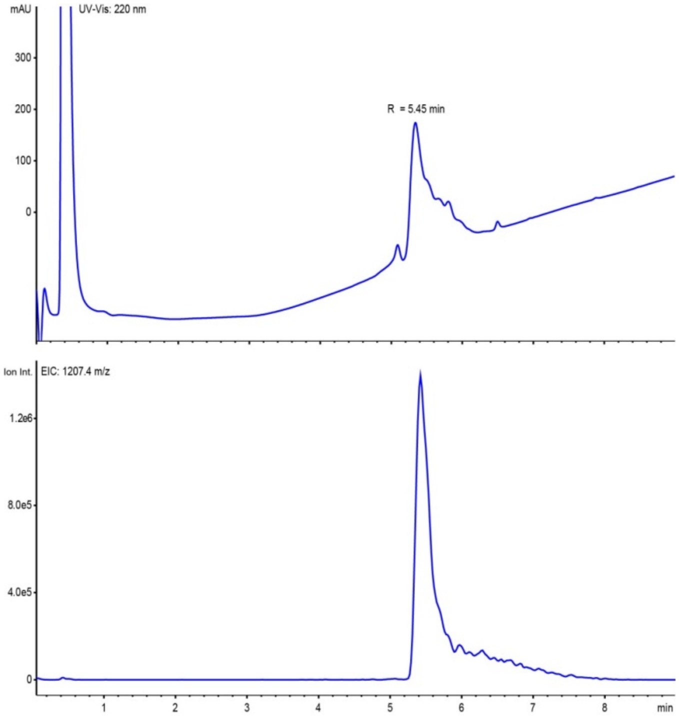
mAU
UV-Vis: 220 nm
R = 5.45 min
300
200
100
0
Ion Int.
EIC: 1207.4 m/z
1.2e6
8.0e5
4.0e5
0
1
2
3
4
5
6
7
8
min

**AND-Gate-FNIR**

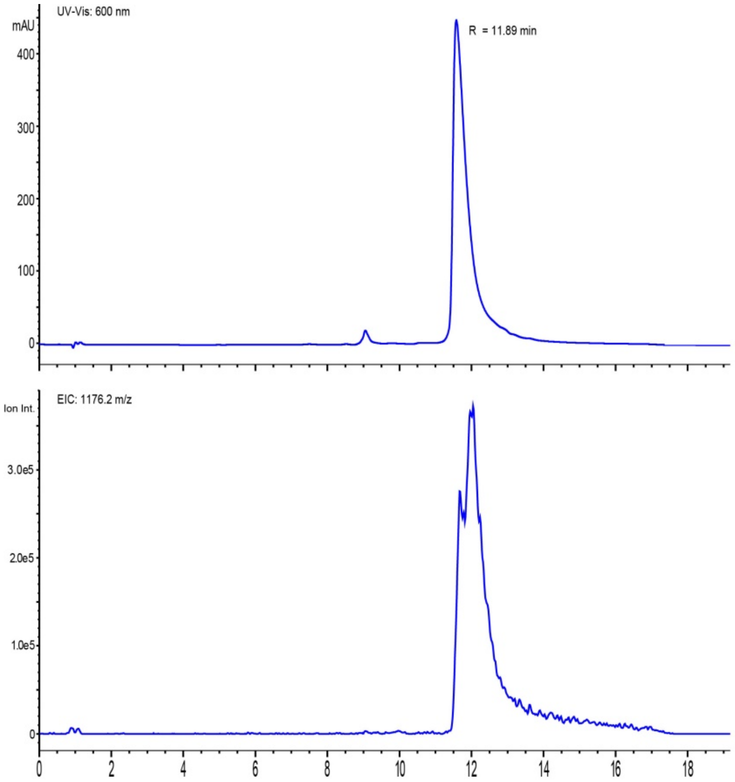
